# TSG101 Associates with PARP1 and is Essential for PARylation and DNA Damage-induced NF-κB Activation

**DOI:** 10.1101/2021.12.07.471537

**Authors:** Ahmet Bugra Tufan, Katina Lazarow, Marina Kolesnichenko, Anje Sporbert, Jens Peter von Kries, Claus Scheidereit

**Affiliations:** Max Delbrück Center for Molecular Medicine in the Helmholtz Association (MDC), Laboratory for Signal Transduction in Tumor Cells, Robert-Rössle-Strasse 10, 13125 Berlin, Germany; Leibniz-Forschungsinstitut for Molecular Pharmacology (FMP), Campus Berlin-Buch, 13125 Berlin, Germany; Max Delbrück Center for Molecular Medicine in the Helmholtz Association (MDC), Advanced Light Microscopy Technology Platform, Robert-Rössle-Strasse 10, 13125 Berlin, Germany

**Keywords:** ADP-ribosylation, DNA damage, IKK-NF-κB pathway, TSG101, breast cancer

## Abstract

In a genome-wide screening for components of the dsDNA-break-induced IKK-NF-κB pathway we identified scores of regulators, including tumor susceptibility protein TSG101. TSG101 is essential for DNA damage-induced formation of cellular poly(ADP-ribose) (PAR). TSG101 directly binds to PARP1 and is required for PARP1 activation. This function of TSG101 is independent of its role in the ESCRT-I endosomal sorting complex. In the absence of TSG101, the PAR-dependent formation of a nuclear PARP1-IKKγ signalosome, which triggers IKK activation, is impaired. According to its requirement for PARP1 and NF-κB activation, TSG101 deficient cells are defective in DNA repair and apoptosis protection. Loss of TSG101 results in PARP1 trapping at damage sites and mimics the effect of pharmacological PARP-inhibition. We also show that loss of TSG101 in connection with inactivated tumor suppressors BRCA1/2 in breast cancer cells is lethal. Our results imply TSG101 as a therapeutic target to achieve synthetic lethality in cancer treatment.

## INTRODUCTION

Endogenous and exogenous sources of DNA damage compromise the genomic integrity of organisms (Jackson and Bartek, 2009). To prevent genome instability or tumorigenesis, cells developed a highly conserved DNA damage response (DDR) that coordinates a network of signaling pathways required for sensing and repairing damaged DNA (Goldstein and Kastan, 2015). Several effector proteins and post-translational modifications (PTM) tightly regulate the DDR (Polo and Jackson, 2011). (ADP-ribose) modification is one of the most crucial regulatory PTMs found in eukaryotes (Gibson and Kraus, 2012). The poly(ADP-ribose) polymerase (PARP) family of enzymes catalyzes ADP-ribosylation by using NAD^+^ as a substrate (D’Amours et al., 1999) and generates the highly negatively charged poly (ADP-ribose) (PAR) polymer (Luo and Kraus, 2012). Poly(ADP-ribose) polymerase 1 (PARP1) is the principal PARP that is involved in DNA damage signaling and produces most cellular PAR (Kim et al., 2005). PARP1 is activated by binding to DNA strand breaks and catalyzes PARylation of target proteins, including itself, histones, and chromatin-associated proteins (Mortusewicz et al., 2007). Enrichment of PAR near the DNA damage site serves as a docking platform for the recruitment of DNA repair factors (Krishnakumar and Kraus, 2010). PARP1 auto-PARylation inhibits its DNA binding capacity and leads to its release from the DNA lesions into the nucleoplasm where it initiates further downstream signaling events (Ahel et al., 2009).

As part of the PARP1 mediated genotoxic stress response, cells activate the NF-κB pathway (Stilmann et al., 2009). Following activation, auto-modified PARP1 forms a transient nuclear complex with IKKγ, PIASy, and ATM (Stilmann et al., 2009). IKKγ is sequentially SUMOylated and phosphorylated in the signalosome complex by PIASy-and ATM (Huang et al., 2003; Mabb et al., 2006; Stilmann et al., 2009). Shortly after the IKKγ modifications, both ATM and IKKγ are exported to the cytoplasm (Wu et al., 2006). ATM triggers autoubiquitination of TRAF6, which recruits cIAP1 and TAB2-TAK1, leading to TAK1 dependent activation of IKKβ (Hinz et al., 2010). After IKK activation, the downstream signaling events occur similar to the classical NF-κB cascade, in which phosphorylation of IκBα by IKK is followed by its ubiquitination and proteasomal degradation, permitting liberated NF-κB dimers to translocate to the nucleus (Hinz et al., 2010; Wang et al., 2017).

The NF-κB pathway upregulates the expression of anti-apoptotic genes, including *Bcl-XL* (Stilmann et al., 2009). Although DNA damage-induced NF-κB has crucial physiological roles, important steps in this signaling cascade are still unexplored. Identification of essential regulators of the DNA damage-induced IKK/NF-ĸB pathway is the basis for the development of directed cancer therapeutics.

Here, we systemically identified regulators of the genotoxic stress-induced NF-ĸB pathway by using genome-wide siRNA screens. Among numerous candidate regulators, we have identified Tumor Susceptibility Gene 101 (TSG101) as an essential component of the DNA damage-induced NF-κB pathway and of PARP1 function in the DNA damage response in general. TSG101 is a member of the endosomal sorting complexes required for the transport complex I (ESCRT-I) (Ferraiuolo et al., 2020); however, its role in the DDR was unexpected. Via biochemical pathway mapping of TSG101 in the IKK-NF-κB signaling cascade, we discovered that TSG101 is essential for enzymatic activation of PARP1. TSG101 interacts through its coiled-coil domain with PARP1 and stimulates PARylation *in vitro* and in intact cells. Depletion of TSG101 completely abrogates cellular PARylation and causes trapping of PARP1 in DNA lesions due to loss of its auto-PARylation. Because of failed PARylation of PARP1, unrepaired DNA foci accumulate. TSG101 is required for the formation of the PAR-dependent DNA damage-induced PARP1-IKKγ signalosome complex. Targeting the TSG101-PARP1 axis sensitized cells to apoptosis due to impaired anti-apoptotic NF-κB driven gene expression. We also demonstrate a co-dependency of TSG101 and BRCA1/2 for the survival of breast cancer cells, similar to the synthetic lethality observed for PARP1 inhibitor olaparib and BRCA1/2 deficiency.

## RESULTS

### Systematic Identification of Essential Components and Regulators of the DNA Damage-induced NF-κB Pathway via Genome-wide siRNA Screens

For a systematical identification of regulators required for the DNA damage-induced NF-κB pathway, we performed a high-content genome-wide siRNA screen. We used a transcription activity-based luciferase assay as a readout for stimulus-dependent NF-κB pathway activation (Fig 1A). After delivery of the siRNA library, etoposide served as DNA damage-inducing agent to activate NF-κB. Cells transfected with *IKBKG* siRNA, which abolished NF-κB-driven luciferase activity, served as positive controls. We selected the 1.000 top hits as candidate activators of the pathway and 100 further hits, which, on the contrary, suppressed NF-κB activation (Fig 1B, Table EV1). The putative positive regulatory hits covered many previously described members and regulators of the NF-κB pathway (Fig 1C, Table EV1). For example, RELA, a principal transactivating NF-κB subunit, topped the list. Moreover, top hits included ATM, TIFA, CHUK, TRAF6, and IKKγ, all known components of the DNA damage-induced NF-κB pathway (Fig 1C). The other side of the screening curve revealed several previously described negative regulators, whose depletion unleashed NF-κB activation (Fig 1B), including NFKBIA, CYLD, TRAF3, VPS28, and N4BP1. Furthermore, we identified TANK, and SENP1, suppressors specifically of genotoxic stress-induced IKK/NF-κB signaling (see Fig 1C for details). Poly(ADP-ribose) glycohydrolase (PARG) is also classified as a negative regulator. Overall, the screening assay revealed a high degree of robustness (Z’>0.5, Table EV2) and technical reproducibility (Fig EV1C.)

**Figure 1.**
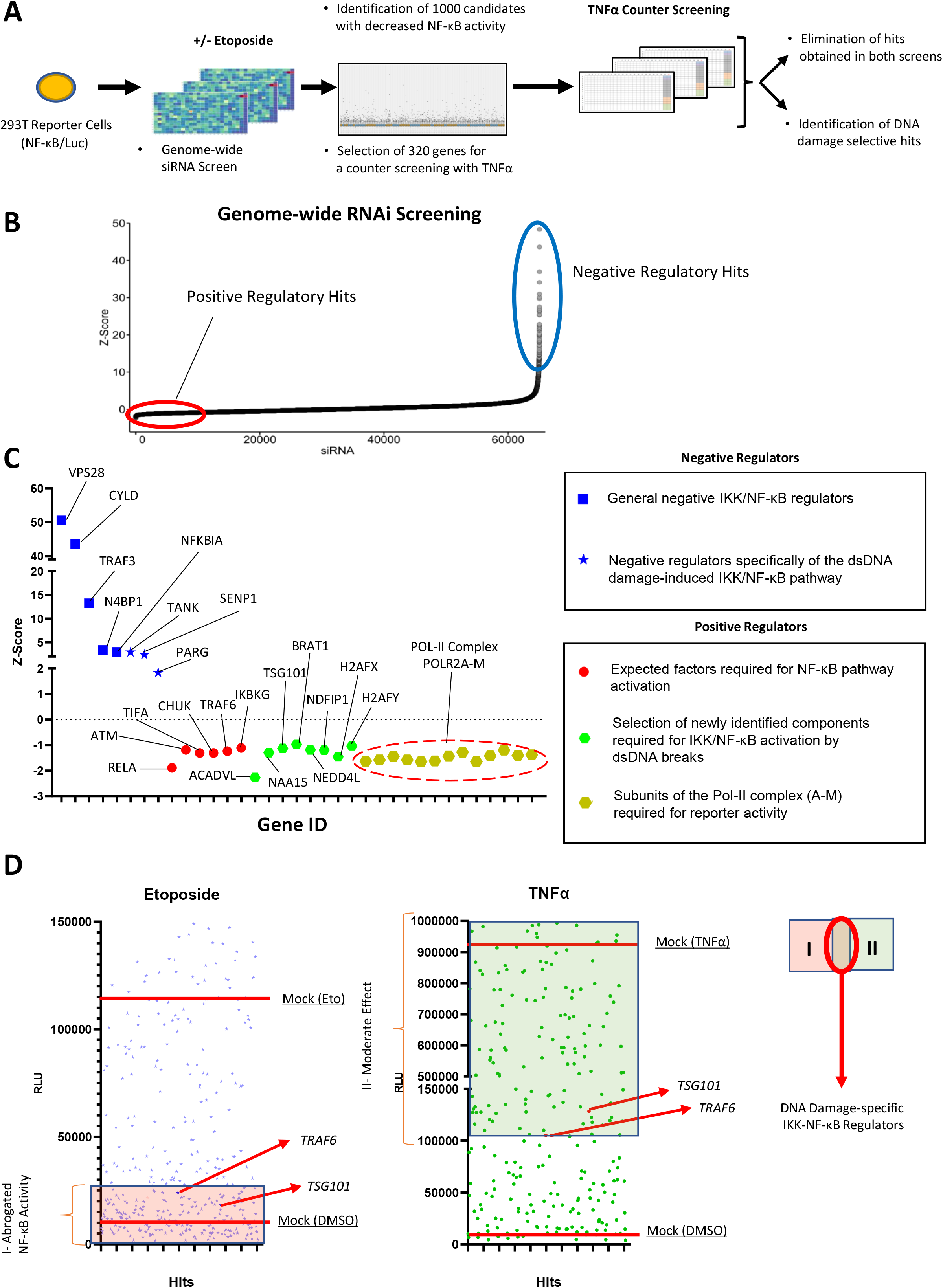
Genome-wide siRNA Screening. A: Schematic diagram of the two consecutive siRNA screens. NF-κB-driven luciferase activity served as a readout for pathway activation. The primary screen was performed using a genome-wide siRNA library. 320 candidates were selected for a secondary counter screen with TNFα to narrow down DNA damage-specific regulators of the NF-κB pathway. Responsiveness of the reporter cell line to NF-κB activating stimuli was analyzed by western blotting (Fig EV1A). B: Cumulative distribution of Z-scores of the genome-wide siRNA screening. Candidates with low Z-scores are putative positive regulators as their depletion abrogated NF-κB activation by DNA damage (red circle). Putative negative regulators with high Z-scores exhibited elevated NF-κB activity (blue circle). C: Z-scores of the best-scoring siRNAs for selected previously known and newly identified pathway regulators. Hits include VPS28 (Mamińska et al., 2016), CYLD (Trompouki et al., 2003), NFKBIA (Haskill et al., 1991), TRAF3 (Vallabhapurapu et al., 2008), N4BP1 (Shi et al., 2021), TANK (Wang et al., 2015), SENP1 (Lee et al., 2011; Shao et al., 2015), PARG (Cortes et al., 2004), RELA (Li et al., 2001), TIFA (Fu et al., 2018), CHUK (Colomer et al., 2019), TRAF6 (Hinz et al., 2010) and IKBKG (Mabb et al., 2006) for expected factors. D: Representative relative luciferase units (RLU) of the differential counter screen of the 320 pre-selected hits are shown. The RLU of hits are displayed for etoposide and TNFα screens (left and right panels). RLU levels of treatment controls are indicated with red lines. Hits that abrogated etoposide-induced NF-κB activity were classified as group I. Hits in the TNFα counter screen with higher activity than in si*TRAF6*-transfected cells constituted group II. The intersection of the two groups indicates putative DNA damage-selective positive regulators. Note that TNFα may trigger secondary NF-κB activation events through autocrine/paracrine mechanisms. See Table EV5 for luminescence values.

We further refined the group of 1000 candidates to 320 (Table EV3), by excluding genes encoding members of general transcription or RNA processing machineries, the proteasome, and ribosome, which are generally required for NF-κB-dependent gene expression. The selected 320 hits were subjected to a counter screen with TNFα as inducer. Targeting of known components of the DNA damage-induced IKK-NF-κB pathway, such as *TRAF6*, which have no documented prominent role in TNFα-induced NF-κB signaling, was included to establish an arbitrary threshold (Fig 1D). Following the counter screen, 60 hits remained, which were highly selective for the DNA damage-induced pathway (Table EV4). To determine functional protein networks, these hits were analyzed in ENRICHR (Kuleshov et al., 2016), REACTOME (Jassal et al., 2020), and STRING (Szklarczyk et al., 2019) databases. The curated networks revealed that the multifunctional tumor susceptibility gene product TSG101 (Ferraiuolo et al., 2020) clusters with several other hits from the primary screen (Fig EV1D), implying a crucial biological function of TSG101 for DNA damage-induced NF-κB signaling.

The candidate hits also included NAA15, an auxiliary subunit of the N-terminal acetyltransferase A (Nat A) complex 15 (Fig EV1E) (Arnesen et al., 2005) (Fig 1C), and several mitochondrial enzymes.

### TSG101 is Essential for the DNA Damage-induced NF-κB Pathway

To validate the contribution of TSG101 to the DNA damage-induced IKK-NF-κB pathway, we knocked down *TSG101* and, as a positive control, *ATM* (Piret et al., 1999). Indeed, depletion of TSG101 or ATM equivalently abrogated NF-κB-driven reporter gene activation (Fig 2A). Furthermore, IKK-dependent phosphorylation of p65 at serine 536, a key step in NF-κB activation after DNA damage (Kolesnichenko et al., 2021), was abolished in TSG101 or ATM deficient cells (Fig 2B). Importantly, the *TSG101* knockdown did not affect activation or protein level of ATM (Fig 2B).

**Figure 2.**
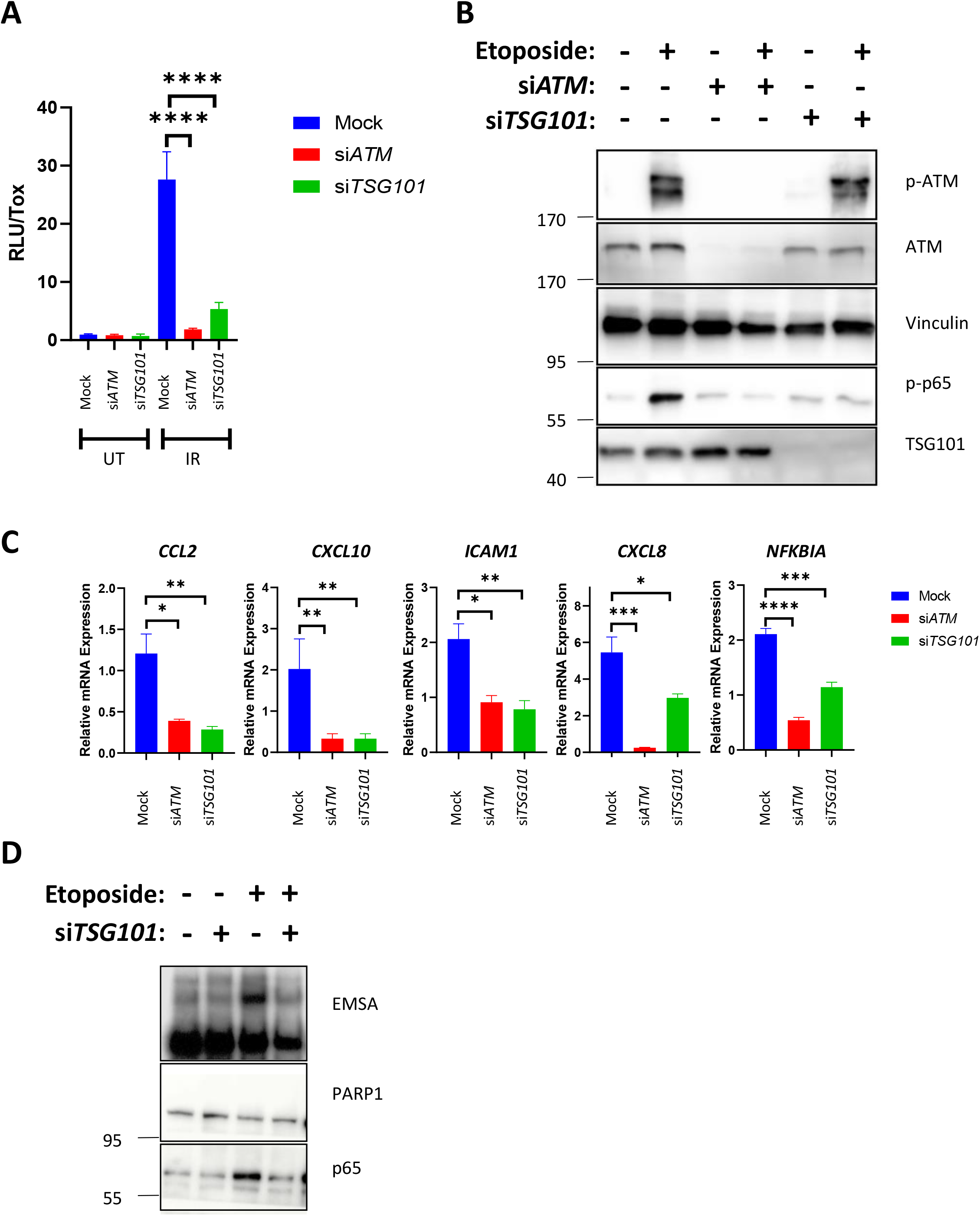
TSG101 is Essential for the DNA damage-induced NF-κB Pathway. A: IR-induced NF-ĸB pathway activation was analyzed via luciferase activity of HEK-Luc cells, transfected with the indicated siRNAs. The indicated cells were irradiated (20 Gy, 8 hours before analysis). A late time point was used for efficient luciferase expression. Luciferase activity was normalized to the viable cell number (TOX fluorescence). The data represent three biological and five technical replicates. For knockdown efficiency, see Figure EV2A. B: U2-OS cells were transfected with the indicated siRNAs. The indicated cells were treated with etoposide (50 µM, 90 min. before analysis). Whole-cell extracts were immunoblotted with the indicated antibodies. The result is representative of three biological replicates. C: U2-OS cells were transfected with the indicated siRNAs and exposed to irradiation (20 Gy, 90 min. before analysis). The expression of indicated genes was analyzed using RT-qPCR. The mRNA expression was normalized to the expression of the housekeeping genes *ACTA1, RPL13A*, and *TBP2*. Data is representative of three biologically independent experiments +/- SEM. D: U2-OS cells were transfected with non-targeting or *TSG101*-targeting siRNAs and irradiated (20 Gy, 90 min. before analysis). Nuclear fractions were analyzed by EMSA (top panel) or immunoblotted with the indicated antibodies (bottom panels).

TSG101 is required for the induced expression of selected NF-κB target genes (*CCL2*, *CXCL10*, *ICAM1*, *CXCL8*, and *NFKBIA*) following irradiation (Fig 2C) or etoposide treatment (Fig EV2B), in the same manner as ATM. In agreement with these results, the knockdown of *TSG101* blocked genotoxic stress-induced nuclear translocation of p65 and DNA binding activity of NF-κB (Fig 2D). Taken together, our data demonstrate that TSG101 is essential for the activation of the NF-κB pathway by DNA damage.

### TSG101 is Required for PARylation

The formation of a transient nuclear PARP1 signalosome complex involving ATM, IKKγ, and PIASy (Stilmann et al., 2009) and a cytoplasmic ATM-TRAF6 module, where ATM is exported to the cytoplasm (Hinz et al., 2010) are cornerstones of the DNA damage-induced IKK pathway. We next analyzed how and where TSG101 acts on this pathway. To investigate if TSG101 affects ATM export, we prepared nuclear and cytoplasmic fractions. In cells treated with etoposide, loss of TSG101 influenced neither activation of nuclear ATM, nor cytoplasmic export of phosphorylated ATM (Fig 3A). Likewise, pharmacological inhibition of PARP1 did not impair ATM export. These data indicate that TSG101 does not function upstream of the cytoplasmic export of phosphorylated ATM.

**Figure 3.**
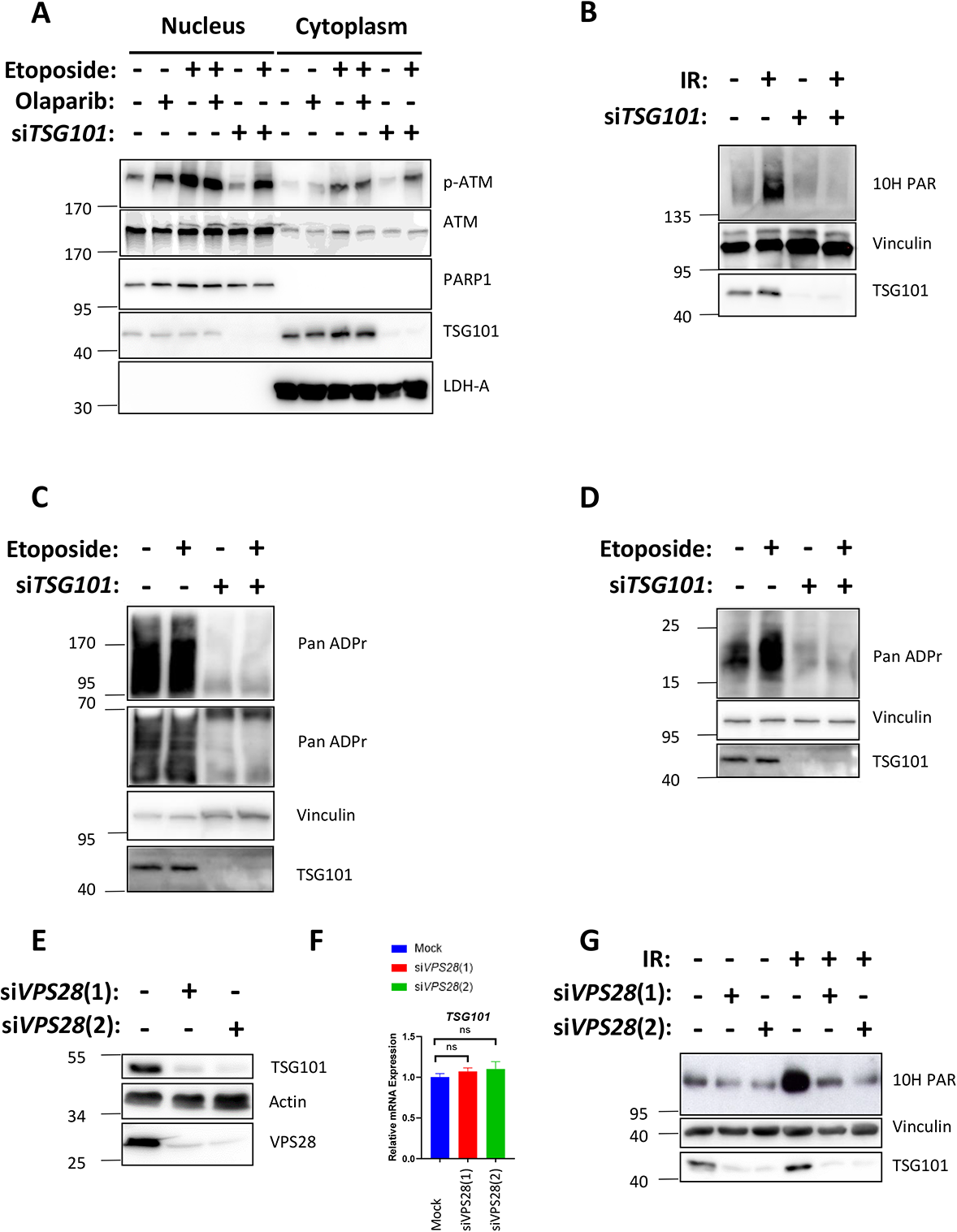
TSG101 Controls PARylation. A: Nuclear-cytoplasmic fractionation of U2-OS cells. Indicated samples were transfected with siRNAs against *TSG101*. PARylation was inhibited by olaparib (10 µM, 24 hours before analysis). Cytoplasmic export of p-ATM was induced by etoposide (50 µM, 45 minutes before analysis). Nuclear and cytoplasmic fractions were immunoblotted with the indicated antibodies. The fractionation efficiency was controlled by the respective subcellular marker proteins (PARP1 and LDH-A). B: U2-OS cells were transfected with non-targeting or *TSG101*-targeting siRNAs. C Cells were irradiated as indicated (20 Gy, 10 min. before analysis). Whole-cell extracts were immunoblotted with antibodies as indicated. Note that the PARylation activated during cell lysis (Jungmichel et al., 2013) was blocked by adding olaparib to the lysis buffer. C: Whole-cell extracts were obtained as in B and immunoblotted with PAN ADPr reagent that selectively recognizes both poly-and mono-ADP-ribose. Two separate gels were used. PARylation during cell lysis was not blocked by olaparib addition to the extraction buffer. D: To analyze ADP-ribosylation of histones, the 25 to 15 kDa range of proteins from C was immunoblotted with the PAN ADPr reagent or the indicated antibodies. E: U2-OS cells were transfected with non-targeting or *VPS28*-targeting siRNAs. Whole-cell extracts were immunoblotted with indicated antibodies. F: Total mRNA was extracted from the samples in E. Expression of *TSG101* was analyzed by RT-qPCR and normalized to two housekeeping genes, *ACTA1* and *RPL13A*. G: Cells were irradiated as indicated (20 Gy, 10 min. before analysis). Whole-cell extracts were immunoblotted with the indicated antibodies. The results are each presentative for three (panels A-D and G) or six (panel F) independent experiments.

The DNA damage-induced formation of the PARP1 signalosome is highly dependent on PARP1 auto-PARylation, which mediates the interaction between PARP1, ATM, IKKγ, and PIASy (Stilmann et al., 2009). Strikingly, in the absence of TSG101, irradiation-induced PARylation was completely abrogated (Fig 3B), implying that TSG101 regulates the DNA damage-induced NF-κB pathway by enabling the formation of the transient PARP1 signalosome complex. Furthermore no poly or mono ADP ribose adducts were formed in TSG101-depleted cells (Fig 3C). In agreement with these observations, both basal level and DNA damage-induced PARylation of histones were severely diminished in the absence of TSG101 (Fig 3D). Depletion of TSG101 does not have any effect on cellular NADH levels in non-irradiated cells (Figure EV3A), excluding that any remote mechanisms control NADH/NAD+ availability needed for PARylation were not altered. Upon irradiation, however, depletion of TSG101 or addition of olaparib decreases the NAD+/NADH ratio, while ectopic TSG101 expression causes an increase (Figure EV3B).

As a member of the ESCRT-I complex, depletion of TSG101 destabilizes the early endosome morphology and dynamics at a wide range (Doyotte et al., 2005). Indeed, it was previously shown that Vacuolar Protein Sorting 28 (VPS28), another member of the ESCRT-I complex (Babst et al., 2000), is diminished in cells lacking TSG101 (Doyotte et al., 2005). Strikingly, we found that TSG101 protein was diminished in VPS28 siRNA-transfected cells (Fig 3E), without affecting the TSG101 mRNA level (Fig 3F), indicating mutual protein stability regulation. Consistent with our previous data, irradiation-induced PARylation was impaired upon VPS28 depletion-mediated reduction of TSG101 levels (Fig 3G). Collectively, our data show therefore that TSG101 functions like an essential cofactor for PARP1 and regulates the DNA damage-induced NF-κB pathway activation by enabling PAR-dependent interactions between ATM, PARP1, IKKγ, and PIASy. Furthermore, the siRNA-mediated depletion of two other newly identified hits, *ACADVL* and *NAA15* (Fig 1C), also blocked cellular PAR formation (Figs EV3C and EV3D). The positive regulatory role of *ACADVL* and *NAA15* in DNA damage-induced NF-κB activation was confirmed by the analysis of etoposide-induced IκBα (Ser 32/36) and p65 (Ser 536) phosphorylation (Figs EV3E and EV3F). A recent study revealed that the NatA complex with its NAA15 and NAA10 regulatory and catalytic components accounts for N-terminal acetylation (Nt-Ac) of a majority of proteins, including PARP1 and TSG101, and that Nt-Ac may play a general role in protein stabilization (Mueller et al., 2021). Indeed, we observed significant decreases in protein levels of PARP1 and TSG101 in NAA15-depleted cells, which may mechanistically explain the reduction in PARylation (Fig EV3G). Altogether, our data reveal several unrelated factors that are required for PARylation and for genotoxic stress-induced IKK/NF-κB activation, underscoring the importance of this post-translational modification for this pathway.

### TSG101 Interacts with and Activates PARP1

To investigate how TSG101 may control PARylation we analyzed physical interactions between TSG101 and PARP1. As a primary notice, TSG101 was predicted to interact with PARP1 and several of the members of the PARP1 signalosome complex (Fig EV4A). We have also previously observed TSG101 as an irradiation-enhanced, likely PARP1-mediated interaction partner of IKKγ in a proteome-wide SILAC-based IP analysis (Mikuda et al., 2018). With a proximity ligation assay (PLA) we were able to show that the interaction of endogenous TSG101 with PARP1 is direct and takes place in the nucleus (Fig 4A). The complex formation took place both, in unstimulated cells and following DNA damage induction by etoposide and was independent of PARylation, as it was insensitive to the PARP inhibitor olaparib (Fig 4A). We also observed that ectopically expressed TSG101 interacts with PARP1 (Fig EV4B).

**Figure 4.**
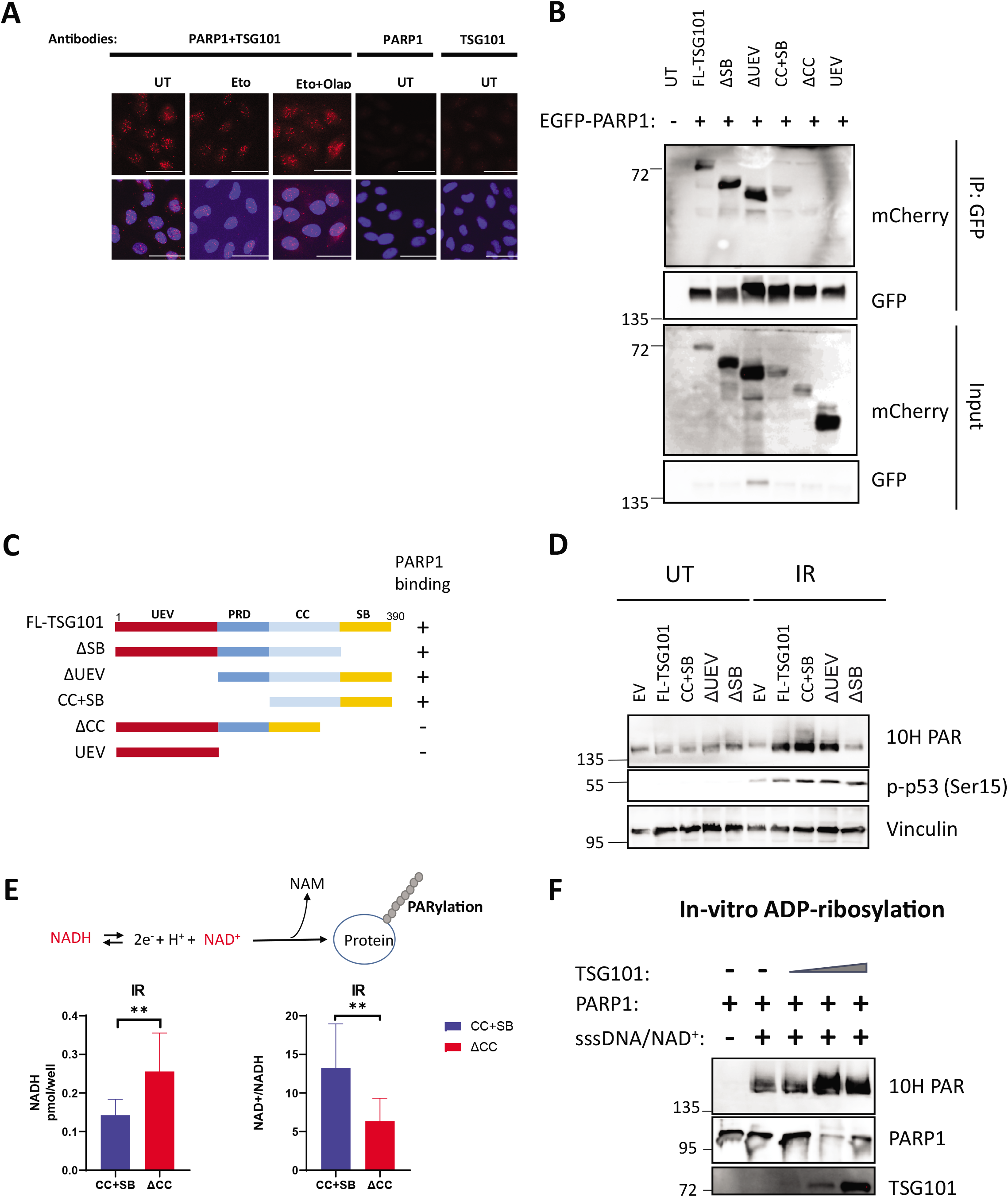
TSG101 Interacts with PARP1. A: Proximity ligation assay (PLA) assay with unstimulated, etoposide (Eto)-treated (50 µM, 10 min. before analysis), or etoposide plus olaparib-(Olap) co-treated (50 µM of etoposide, 10 min. before analysis and 10 µM of olaparib 24 hours before analysis) U2-OS cells. The red dots represent PARP1-TSG101 interaction. The nucleus is stained with DAPI (blue). Scale bar: 30 µm. As negative controls, PLA was performed without primary PARP1 or TSG101 antibodies. The data in A-C are each representative of three biologically independent experiments. B: GFP-tagged PARP1 and mCherry-tagged full-length and truncated versions of TSG101 were ectopically expressed in U2-OS cells. GFP was immunoprecipitated from the whole-cell extracts and samples were immunoblotted as indicated. C: Summary of results of Fig 4B. PARP1 interacting constructs were indicated with a “+”. D: U2-OS cells were transfected with the indicated constructs (The Coiled-coil containing CC+SB and the ΔCC). Cells were irradiated (20 Gy, 5 minutes before analysis) and the decomposed NADH and total NAD levels were quantified. NADH/NAD^+^ is shown in the right panel. E: *In-vitro* PARylation reactions were carried out with the indicated recombinant and purified proteins. Increasing amounts of TSG101 (25, 50 and 100 ng) were added to PARP1 (8 ng), as indicated. sssDNA: sheared salmon sperm DNA.

We then determined the minimal region of TSG101 necessary for PARP1 interaction and found that the coiled-coil (CC) domain of TSG101 is essential for its association with PARP1 (Figs 4B and C). Strikingly, ectopic overexpression of full-length TSG101 or its PARP1-interacting truncated versions enhanced DNA damage-induced PARylation (Fig 4D). To further prove that the TSG101 CC domain enhances the catalytic activity of PARP1, we tested the effect of this domain on irradiation-induced consumption rates of NADH (Fig 4E). In agreement with our previous observation, expression of the CC+SB domains of TSG101 but not the CC-deleted TSG101 version significantly enhanced the NADH consumption (Fig 4E). To examine more directly the effect of TSG101 on the catalytic activity of PARP1, we performed an *in vitro* PARylation assay. As expected, PARylation was catalyzed upon incubation of recombinant PARP1 with NAD^+^, MgCl_2_, and sheared salmon sperm DNA (Fig 4F). Strikingly, addition of TSG101 enhanced the catalytic activity of PARP1 in a dose-dependent manner (Fig 4F). Collectively, our data suggests that the interaction between PARP1 and TSG101 is direct and functionally important for PARylation.

### TSG101 Prevents Trapping of PARP1 in DNA lesions

Rapid recruitment of PARP1 to the DNA lesions enables its direct interaction with the DNA breaks, which triggers its enzymatic activity (Mortusewicz et al., 2007). Therefore, TSG101 might be required either for the recruitment of PARP1 to DNA damage sites or for its dissociation. To gain insights into the mechanism how TSG101 controls these processes, we analyzed the dynamics of PARP1 enrichment at DNA lesions upon laser microirradiation by live-cell imaging (Fig 5A). PARP1 rapidly accumulated at the DNA damage sites in control cells and in cells with TSG101 knockdown (Fig 5B). While PARP1 foci showed dissociation within minutes after micro-irradiation in control cells, it remained captured on DNA lesions in both si*TSG101*-transfected and olaparib-treated cells (Fig 5B). Normal recruitment of PARP1 to DNA lesions in the absence of TSG101 rules out that TSG101 will direct PARP1 to the DNA breaks. A quantitative analysis of the kinetics of PARP1 recruitment revealed that PARP1 association with DNA lesions occurs in the absence of TSG101 in a kinetically similar manner to PARP1 inhibition by olaparib (Figs 5C and 5D). We next investigated if TSG101 would accumulate at DNA damage sites. Unexpectedly, GFP-TSG101 did not migrate to the DNA lesions (data not shown); however, an indirect immunofluorescence analysis of DNA damage sites (Fig 5E) revealed that TSG101 is abundantly present in the nucleus and evenly distributed throughout the nucleoplasm and including the DNA damage sites (Fig 5F). Collectively, our data indicate that due to the requirement of TSG101 for PARylation, PARP1 remains trapped on DNA lesions in its absence.

**Figure 5.**
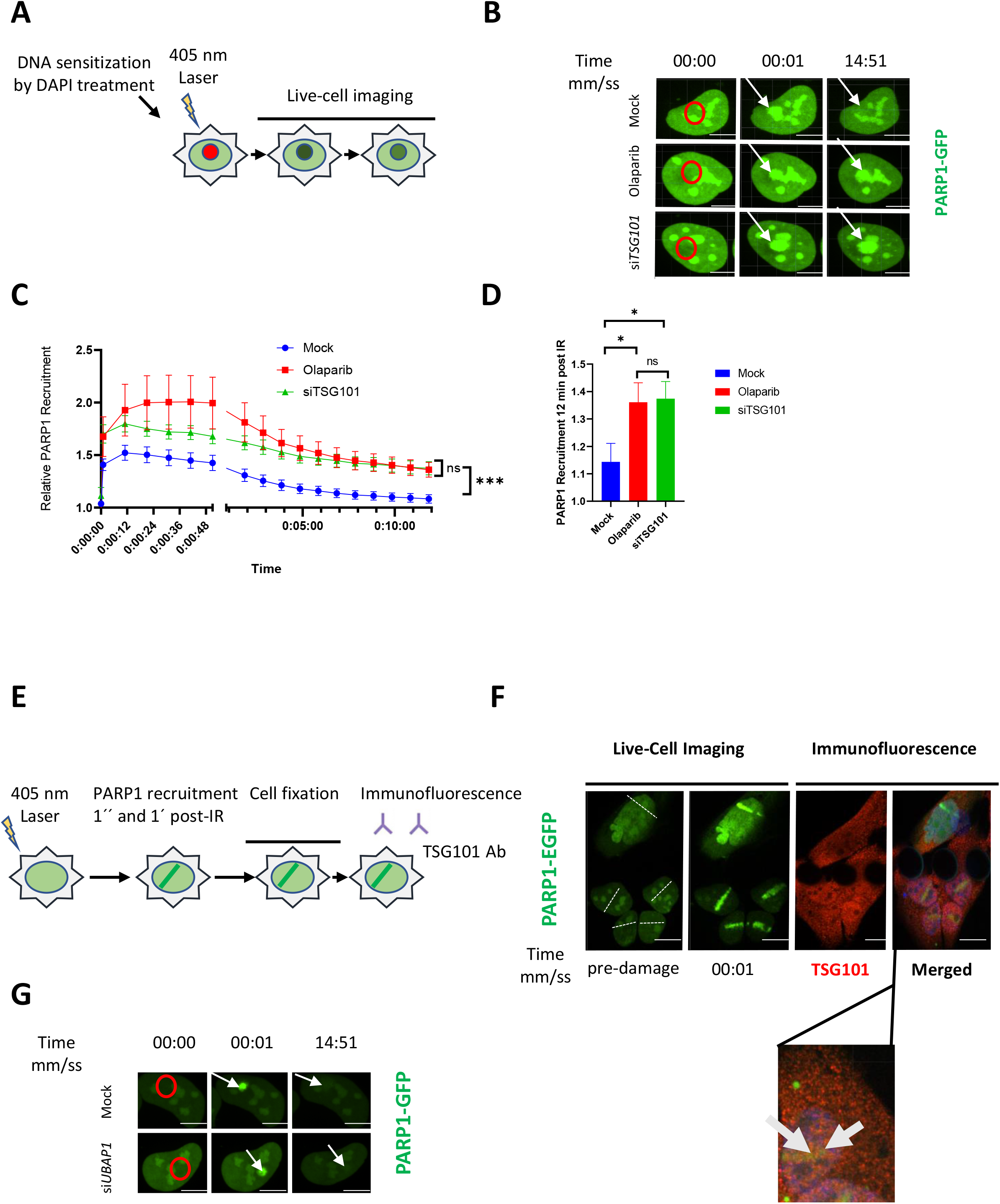
PARP1 is Trapped in DNA Lesions in the Absence of TSG101. A: Schematic of the live-cell PARP1 recruitment assay. U2-OS cells transfected with GFP-tagged PARP1 were sensitized with DAPI (10 µg/ml, for 10 min.). After 405 nm microirradiation, PARP1 recruitment to and dissociation from the DNA lesion was recorded over time. B: Representative images of PARP1-GFP association with laser-microirradiation sites in untreated (mock), olaparib-treated (10 µM, 24 hours before analysis) or si*TSG101*-transfected U2-OS cells at indicated times post-irradiation. Scale bar: 7.5 µm. The image is representative of nine replicates from three biologically independent experiments. Go to supplementary videos for the indicated conditions. C: Kinetics of PARP1-GFP recruitment and dissociation from DNA lesions were measured at times indicated. Nine nuclei were analyzed for each indicated condition from three biologically independent experiments. The data are shown as mean GFP intensity in the microirradiated area ± SEM, normalized to the mean GFP intensity of corresponding whole nuclei. D: Relative PARP1-GFP recruitment to DNA lesions for each indicated condition at 12 min. post-irradiation. E: Scheme of indirect immunofluorescence analysis of DNA damage sites. U2-OS cells were GFP-PARP1-transfected and microirradiated as in A. Cells were fixed shortly after rapid PARP1 recruitment to the DNA lesions (1 min post-microirradiation) and stained for TSG101 by indirect immunofluorescence. F: Indirect immunofluorescence of TSG101 at indicated times post damage. DNA lesions were generated as in B. Stippled lines indicate the applied laser beam. Recruitment of PARP1 to DNA lesions was recorded from live cells. Cells were fixed at 1 min post-microirradiation and stained for TSG101. For specificity of the TSG101 staining see Fig EV5A. G: Representative images of PARP1-GFP association/dissociation at laser-microirradiation sites in untreated (mock) or si*UBAP1*-transfected U2-OS cells at indicated times post-irradiation. Scale bar: 10 µm. The image is representative for four independent experiments.

Based on our screening results we hypothesized that the role of TSG101 in PARylation is independent from its ESCRT-related functions as TSG101 was the sole member of the ESCRT complex that abrogated the DNA damage-induced NF-κB activity (Figure EV5B). We sought to determine whether disruption of the ESCRT machinery affects PARylation. Therefore, we analyzed the recruitment kinetics of PARP1 to the DNA lesions following depletion of UBAP1, another component of the ESCRT-I complex (Ferraiuolo et al., 2020). Importantly, efficient *UBAP1* knockdown did not affect TSG101 protein level (Figures EV5C and D). Dissociation of PARP1 from the laser microirradiated areas occurred normally in si*UBAP1*-transfected cells (Fig 5G), suggesting that the defective ESCRT system does not influence PARylation.

Collectively our data revealed that TSG101’s function in PARylation is independent of its role in the ESCRT complex.

### Loss of TSG101 Sensitizes Cells to Apoptosis and Impairs DNA Repair

Since impaired enzymatic PARP1 activation should result in genomic instability and decreased cell viability, we analyzed cell death and DNA damage in TSG101 depleted cells. Notably, the knockdown of TSG101 severely reduced cell viability following DNA damage in the same manner as an ATM knockdown (Figs 6A and 6B). Furthermore, in the absence of TSG101, DNA damage-induced expression of a crucial anti-apoptotic NF-κB target gene, *Bcl-XL* (Stilmann et al., 2009), was impaired (Fig 6C). Loss of TSG101 also significantly increased the expression of pro-apoptotic genes, including PUMA (Figs EV6A and B).

**Figure 6.**
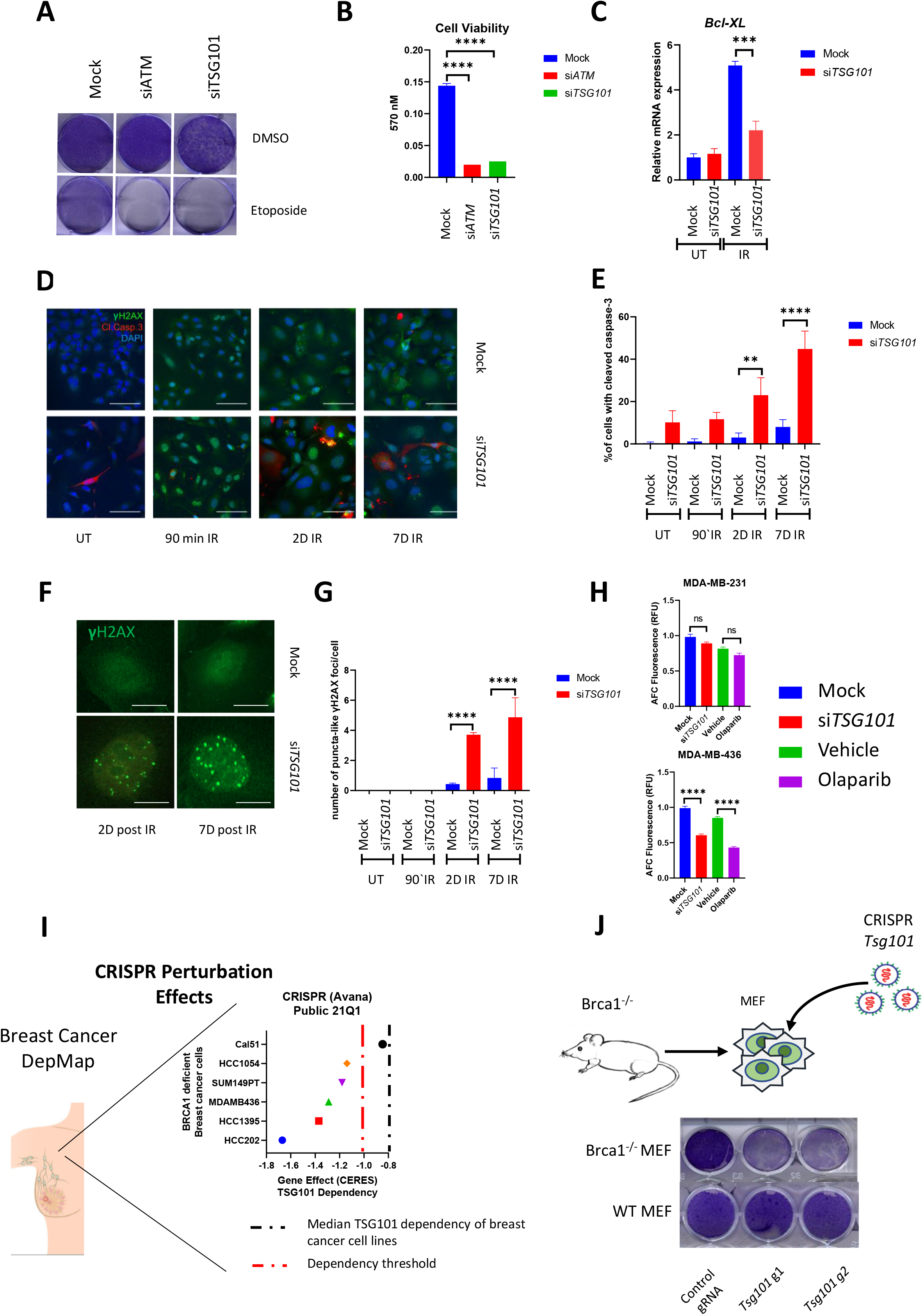
Loss of TSG101 Sensitizes Cells to Apoptosis and Impairs DNA Repair. A: Crystal violet cell viability assay with siRNA-transfected U2-OS cells. Cells were treated with etoposide (50 µM) 72 hours prior to analysis. B: Relative crystal violet intensities of etoposide-treated cells in A were measured at 570 nm. C: U2-OS cells were transfected with TSG101-siRNAs, irradiated (20 Gy) and analyzed 1 day later. Total RNA was extracted and the expression of *BCL2L1* analyzed by RT-qPCR. *BCL2L1* expression was normalized to the housekeeping genes *ACTA1, RPL13A*, and *TBP2*. D: Cells were transfected with non-targeting or *TSG101*-targeting siRNAs. Immunofluorescence staining was performed with the indicated antibodies (red: cleaved caspase 3; green: γH2AX). Nuclei were stained with DAPI (blue). The image is a representative of 3 biologically independent experiments. Scale bar: 30 µm. See Fig EV6D for percentage of γH2AX positive cells as an evidence of IR-treatment. E: Percentage of cleaved caspase 3 was determined by blind counting of approximately 100 cells for each condition from 3 biologically independent experiments. F: γH2AX staining of *TSG101*-targeting or non-targeting siRNA transfected U2-OS cells reveals puncta-like foci at 2 or 7 days after irradiation. Scale bar: 30 µm. G: The number of puncta-like γH2AX foci per cell was determined by blind counting of approximately 100 cells for each condition from 3 biologically independent experiments. H: MDA-MB-231 and MDA-MB-436 cells were transfected with *TSG101*-targeting or non-targeting siRNAs. Cells were treated with vehicle (DMSO) or olaparib (10 µM) 96 hours prior to the analysis, as indicated. Relative number of viable cells in culture was measured using a fluorogenic, cell-permeant, peptide substrate (glycyl-phenylalanyl-aminofluorocoumarin; GF-AFC). See Fig EV6G for brightfield images of cells expressing *TSG101*-targeting or non-targeting siRNAs. I: Analysis of breast cancer cell lines bearing damaging *BRCA1/2* mutations in the DepMap database. The CERES score represents the gene effect, which is based on cell depletion assays of the CRISPR (Avana) Public 21Q1 dataset. A CERES score lower than (−1) indicates a high likelihood that the gene of interest is essential in the given cell line. Correspondingly, a - 1 CERES score was indicated with a discontinued red line. The median dependency of all breast cancer cell types to TSG101 expression was indicated with a discontinued black line. J: *BRCA1* deficient MEF cells were lentivirally transduced with two independent guide RNAs targeting mouse *Tsg101* or with empty CRISPR. Cell viability was analyzed with crystal violet staining. CRISPR knock-out of *Tsg101* in wild-type MEF cells did not cause cell death. The knockout efficiency in wild-type cells was determined by immunoblotting (See Fig EV6E).

Induction of apoptosis was visualized by cleaved caspase 3 (Fig 6D). Strikingly, over 50% of the cells stained positive for cleaved caspase 3 at 7 days post-irradiation (Fig 6E). Furthermore, depletion of *Tsg101* in murine cells also increased IR-induced caspase-3 activation (Fig EV6C). Importantly, the early stages of cell death are associated with size increase in γH2Ax foci (Bonner et al., 2008; Solier and Pommier, 2009) and accordingly, puncta-like γH2Ax foci emerged only in irradiated and TSG101-depleted cells at later time points (Figs 6F and 6G).

Altogether, our data suggest targeting TSG101 sensitizes cells to the DNA damage-induced apoptosis. We additionally examined cells at 7 days post-irradiation. As we have recently shown, DNA damage causes a biphasic NF-κB activation, with the second phase occurring days after the initial exposure in a PARP1-and IKK-independent manner and driving the expression of senescence-associated-secretory phenotype (SASP) genes (Kolesnichenko et al., 2021). Therefore, we analyzed *bona fide* SASP factors IL-6 and IL-8 (Fig EV6F). The expression of these SASP factors was significantly upregulated at seven days post-irradiation irrespective of the presence of TSG101. These data underline that TSG101 does not play a role in the regulation of SASP expression.

A clinically implemented synthetic lethality principle is based on the finding that cancer patients carrying BRCA1/2 mutations are sensitive to an inhibition of PARP1 (Helleday, 2011). This prompted us to investigate the effect of TSG101 loss in the context of *BRCA1/2* deficiency. Strikingly, depletion of TSG101 significantly reduced the viability of *BRCA1* mutant cells (MDA-MB-436), whilst *BRCA1* wild type cells (MDA-MB-231) were unaffected (Fig 6H). Furthermore, targeting TSG101 affected cellular integrity of *BRCA1* deficient cells in a similar manner as pharmacological inhibition of PARylation by olaparib (Fong et al., 2009)(Fig 6H). As a second line of evidence, we analyzed the dependency status of *BRCA1/2* deficient breast cancer cells to TSG101 expression in the DepMap database (Tsherniak et al., 2017). In agreement with our findings, five out of six breast cancer cell lines bearing *BRCA1/2* mutations showed a strong dependency on TSG101 (Figure 6I), while the sum of breast cancer cell lines was less sensitive to the loss of TSG101 (Figure 6I), indicating that loss of PARylation in the TSG101 knockout might cause synthetic lethality in these cells. To obtain further evidence, we also targeted *Tsg101* in *Brca1*^-/-^murine embryonic fibroblasts (MEF) with a CRISPR system. A knockout of *Tsg101* in the *Brca1* null background (Fig EV6E) indeed resulted in severely increased cell death (Fig 6J). Thus, in the context of *BRCA1/2* deficiency, TSG101 has a comparable effect on cell survival as a pharmacological PARP1 inhibition. Taken together, our data demonstrate that TSG101 prevents DNA damage-induced cell death by enabling PARylation and subsequent PARP1-dependent NF-κB pathway activation and plays a crucial role in PARP1-dependent DNA repair processes.

## DISCUSSION

### Genome-wide siRNA Screening Revealed Multiple Regulators of the DNA Damage-Induced IKK-NF-κB pathway

NF-κB signaling is important for many physiological processes; however, a genome-wide loss of function screen assessing the role of each particular gene product in the regulation of a specific pathway, to the best of our knowledge, is presented here for the first time. With the advent of high-throughput screening technologies, analysis of the signaling pathway dynamics is now possible, and we applied this approach to systematically identify factors required for regulation of DNA damage-induced IKK-NF-κB signaling. By combining the screening outcome of genotoxic stress-induced signaling with a counter screening using TNFα, we were not only able to identify DNA damage selective regulators of the NF-κB pathway but also provide a comprehensive overview of general regulation of NF-κB signaling. Our dataset underscored previously described key elements of IKK-NF-κB signaling. Importantly, the essential components of the pathway led to abrogated activity in the screen, while targeting negative regulators unleashed the NF-κB pathway (Fig 1D). Previously described negative regulators included TRAF3, which suppresses non-canonical NF-κB signaling by degrading NIK (Sun, 2017), and ESCRT-I components, including VPS28, the depletion of which leads to the accumulation of TNFR receptors in the endosomes, and subsequent activation of NF-κB (Mamińska et al., 2016). More essentially, depletion of PARG, which is an enzyme that recycles the PAR chains and attenuates PARylation (Cortes et al., 2004), also enhanced the DNA damage-induced NF-κB activation. As expected for the nuclear DNA damage initiated and PARP1 dependent cascade (Dunphy et al., 2018), the cGAS-STING pathway did not contribute to NF-κB activation (MB21D1 and TMEM173 in Table EV1). Collectively, these results suggest that our screening results represent a rich resource that provides previously unknown protein complexes or biological processes required for or linked to NF-κB signaling, as well as negative regulators that restrict aberrant activation.

### DNA Damage Induced Activation of the IKK-NF-κB Pathway Highly Depends on PARylation

In response to DNA damage, a transient PARP1 signalosome complex is formed containing the DNA damage sensors ATM and PARP1 bound to IKKγ and PIASy (Stilmann et al., 2009). The interactions between the proteins in this complex are facilitated by PARylation and are requisite for the DNA damage-induced activation of the NF-κB (Stilmann et al., 2009). The strong dependence of the activation of the NF-κB pathway on PARylation enabled us to identify components that regulate PARylation. We provide compelling evidence that three candidate hits, ACADVL, NAA15, and TSG101, are required for PARylation (Figs 3 and EV3). Intriguingly, a recent study revealed that mitochondria-derived NAD^+^ is consumed by nuclear PARP1 in response to DNA damage (Hopp et al., 2021), which supports our ACADVL data, as this enzyme contributes to NAD^+^ homeostasis through fatty acid beta-oxidation (McAndrew et al., 2008). Furthermore, a large group of metabolic enzymes (47 hits), which fall into the same category as ACADVL, was identified in our screening dataset with very high confidence (Table EV6). Investigation of these components will decipher the metabolic networks underlying the NAD homeostasis and NAD-dependent PARylation under genotoxic stress conditions. We also showed that NAA15 is required for stabilization of PARP1 and TSG101 protein levels (Fig EV3G), suggesting that NAA15 may regulate PARylation through controlling Nt-Ac-dependent stability and/or activity of several PAR-regulating factors. The high prevalence of hits encoding genes that are important for PARylation underscores the crucial role of this posttranslational modification for the genotoxic stress-induced IKK-NF-κB pathway.

We also provide evidence that TSG101 enables PARylation through direct protein-protein interaction with PARP1 (Figs 3 and 4). We showed that TSG101 exhibits a robust interaction with PARP1 that occurs independently of PARylation and induction of DNA damage (Fig 4A). In contrast to PARP1, we did not observe an accumulation of TSG101 at the damage sites (Fig 5E). However, TSG101 is present throughout the nucleus and including the DNA lesions and is therefore available for direct interaction with and activation of PARP1.

We have previously identified proteins that associate with increased strength with IKKγ after DNA damage using a SILAC-based IKKγ pull-down (Mikuda et al., 2018). These factors included TSG101, as well as further hits of our genome-wide siRNA screen (Table EV7). The increase of TSG101 binding to IKKγ after DNA damage (Mikuda et al., 2018) can be explained by the much stronger binding of IKKγ to PARylated PARP1 (Stilmann et al., 2009), while TSG101 is drawn into this complex by direct, PAR-independent association with PARP1 (Fig 4A). We further showed that CC domain of TSG101 is crucial for PARP1 interaction (Fig 4B) and ectopic overexpression of only-CC containing versions or the full-length TSG101 enhanced the IR-induced PARylation (Fig 4D). In agreement with these observations, TSG101 hyperactivated PARP1 in an *in vitro* system (Fig 4F). Intriguingly, a recent study showed that TSG101 binds with its CC domain to nuclear glucocorticoid receptor (GR) and folds a disordered region in GR to enhance its transcriptional activity (White et al., 2021). Thus, we speculate that a similar scenario may underlie the enzymatic activation of PARP1. Recent discoveries of PARP1 interaction partners (HPF1 (Gibbs-Seymour et al., 2016), CHDL1 (Blessing et al., 2020), and CARM1 (Genois et al., 2021)) revealed several levels of modulation of the enzymatic activity of PARP1. However, none of these proteins act in a manner functionally equivalent to TSG101, and their depletion did not result in complete loss of cellular PARylation. Future determination of the structure of TSG101 bound to PARP1 will reveal how this interaction mediates PARylation and this may facilitate the development of drugs to modulate PARP1 activity.

### TSG101 Regulates PARylation Independently of the ESCRT Complex

One major issue targeting TSG101 would be its functions in cellular homeostasis, particularly through the ESCRT system. Our screening results suggested that the role of TSG101 in PARylation is independent of its ESCRT function as TSG101 was the sole ESCRT member that abrogated the DNA damage-induced NF-κB activation (Table EV1, Figure EV5B). Importantly, it was previously shown that targeting the ESCRT complex members perturbs TNFR family internalization and upregulates the NF-κB pathway (Mamińska et al., 2016). Depletion of VPS28, UBAP1, and CHMP4B all led to high superactivation, only TSG101 to inhibition of NF-κB activity (Table EV1). There is an overlap of the two functions for TSG101, activator in the DNA damage pathway and negative regulator endocytosis-related NF-κB activation. To further support the finding that TSG101 controls PARylation independently of the ESCRT complex, we demonstrated that the UEV domain of TSG101, which is responsible for binding to ubiquitinated cargo proteins and mediating the ESCRT-related functions (Vietri et al., 2020), does not interact with PARP1 (Fig 4B). Importantly, we showed that TSG101 is found in the nucleus (Fig 5F) at an abundant level, unlike the other ESCRT members, which are localized in the cytoplasm and in endosomes (Thul and Lindskog, 2018), revealing the existence of ESCRT-independent nuclear functions of TSG101. We further demonstrated that another member of the ESCRT complex, UBAP1, does not influence the dissociation of PARP1 from DNA lesions (Fig 5G). Taken together, our data suggests that pharmacological targeting of TSG101-PARP1 complex formation would not interfere with other essential cellular functions of TSG101.

### TSG101 is Crucial for DNA Repair and Synthetically Lethal with BRCA1/2

Previous attempts to generate a *Tsg101*^-/-^ mouse model revealed that its knockout is embryonically lethal (Ruland et al., 2001). Intriguingly, p53 protein accumulation, which was independent of Mdm2 and p53 transcript levels, was observed in the homozygous Tsg101^-/-^ embryos (Ruland et al., 2001). This observation further supported our findings that TSG101 is required for PARylation because the increased p53 signaling in Tsg101 null embryos might be a consequence of prolonged loss of PARylation, which is known to accelerate replication fork speed above a tolerated threshold and initiates DNA damage response (Maya-Mendoza et al., 2018). Indeed, the requirement of TSG101 for PARylation implicates that TSG101 might be indispensable for DNA repair functions. We showed that in the absence of TSG101, γH2AX foci, likely caused by increased apoptosis, appear days after DNA damage generation as puncta-like large foci (Figures 6F and G). In line with this evidence, we showed that *BRCA1/2* deficient breast cancer cell types have a strong dependency on TSG101 as demonstrated by the cell viability scores (Fig 6H) and the DepMap database (Tsherniak et al., 2017) (Fig 6I). Taken together, our results (see Fig 7 for a global summary) implicate that pharmacological targeting of the TSG101-PARylation signaling axis could exploit synthetic lethality in tumors and impede tumor growth.

**Figure 7.**
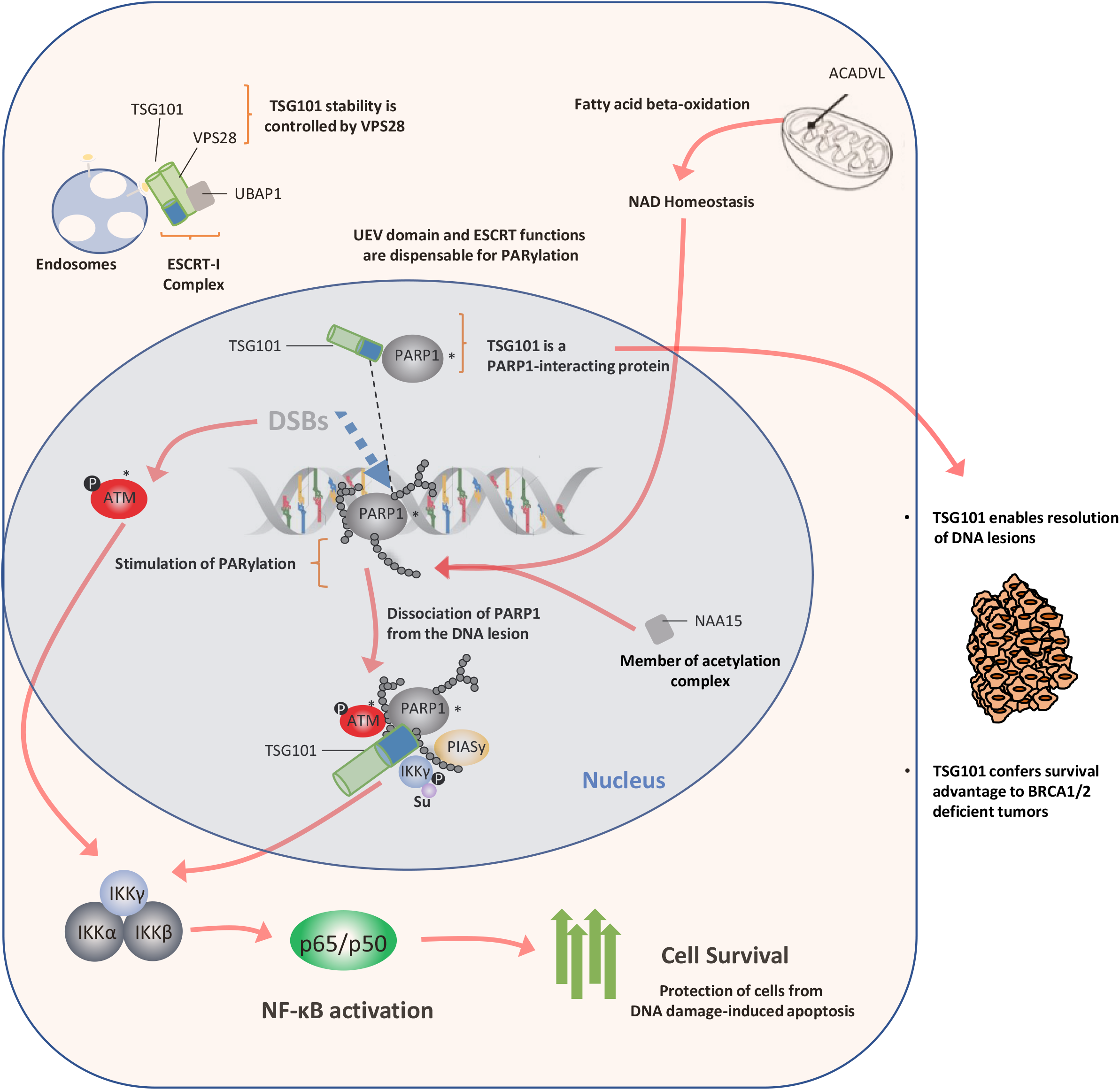
Schematic Summary. Activation of the NF-κB pathway by DNA damage depends on PARylation dependent formation of a transient nuclear PARP1 signalosome complex with ATM, PIASy, and IKKγ. ACADVL, NAA15 and TSG101 are all required for PARylation and PAR-dependent DNA damage-induced NF-κB activation. ACADVL, an acyl-coenzyme A dehydrogenase, may control homeostasis of mitochondrial NAD^+^, known to be required for nuclear PARP1 activation following DNA damage. The N-terminal acetylase regulatory component NAA15 controls the stability or activity of factor(s) needed for PARylation. TSG101 directly interacts with PARP1 and is required for dissociation of activated PARP1 from DNA lesions. TSG101-dependent PARylation and formation of the TSG101-PARP1 complex is independent of the role of TSG101 in the ESCRT complex. Activation of NF-κB through the TSG101-PARylation axis protects cells from DNA damage-induced apoptosis. TSG101 is also crucial for DNA repair functions and is synthetically lethal in conjunction with BRCA1/2 mutations.

## ACKNOWLEDGEMENTS

We thank Dr. Michela Di Virgilio for discussion and Brca1^-/-^ MEF cells. A.B.T. acknowledges DFG, GSC 1091, Berlin Graduate School for Integrative Oncology (B.S.I.O.), and Deutscher Akademischer Austauschdienst (DAAD) for support. The work was funded in part by grants from BMBF, CancerSys project ProSiTu and Helmholtz Association iMed to C.S.

## AUTHOR CONTRIBUTIONS

A.B.T. optimized the reporter cell system for screening. K.L. performed the siRNA library screening and primary screening data acquisition. A.B.T. carried out all functional analyses of hits. A.B.T. and A.S. performed microirradiation microscopy and data analysis. M.K. performed H2AX/cleaved caspase-3 staining and EMSA. J.P.v.K. and C.S. conceived the screening strategy. A.B.T. and C.S. conceived the experimental plan and prepared the manuscript with input from all authors.

## MATERIALS AND METHODS

### Cell Lines

All cell lines were grown under sterile and standard cell culture conditions (humidified atmosphere, 5% CO_2_) and routinely tested for mycoplasma.

The NF-κB/293/GFP-Luc^TM^ cell line was cultured with Roswell Park Memorial Institute Medium (RPMI) supplemented with 10% FBS.

293T, MEF, and U2-OS cell lines were cultured in high glucose-containing Dulbecco’s modified Eagle’s medium (DMEM) supplemented with 5% penicillin/streptomycin (P/S) and 10% FBS. The *BRCA1*^-/-^ MEF cells (Callen et al., 2013) were kindly provided by Dr. Michela Di Virgilio.

The TSG101 CRISPR knock-out bulk cells of U2-OS and MEF origin were cultured in high glucose-containing Dulbecco’s modified Eagle’s medium (DMEM) supplemented with 5% penicillin/streptomycin (P/S), 10% FBS, and 2µgr/ml puromycin.

MDA-MB-231 cells were cultured in Dulbecco’s modified Eagle’s medium (DMEM) supplemented with 5% penicillin/streptomycin (P/S), 10% FBS and 5% non-essential amino acids. MDA-MB-436 cells were cultured in RPMI 1640 medium supplemented with 5% penicillin/streptomycin and 20% FBS.

### High-content siRNA Screening

The screening was performed with a genome-wide siRNA library (Ambion Silencer^R^ Human Genome siRNA library v3, Thermo Fisher), composed of siRNAs targeting approximately 21000 genes arrayed on 384-well plates. Each screening plate contained several controls such as a non-targeting siRNA (Silencer Negative Control from Ambion), a cell-death inducing siRNA mixture, (AllStars Hs Cell Death Control siRNA from Qiagen), and siRNA against IKKγ. For a detailed plate layout see supplementary Figure 1B.

The NF-κB-driven luciferase activity of the reporter cells (NF-κB/293/GFP-Luc^TM^ cell line) was used as a pathway readout. For the transfection of the siRNA library, on a Freedom EVO 200 workstation (Tecan) 5 µl of a 500 nM library-siRNA-OptiMEM solution was mixed in each well of the 384-well assay-plate with 0.05 µl Lipofectamine RNAiMAX transfection reagent previously diluted in 4.95 µl OptiMEM (Life Technologies). Subsequently, 1000 cells/well in 40 µl antibiotic-free complete medium were seeded onto the pre-dispensed transfection mixture using an EL406^TM^ (BioTek) dispenser resulting in a 50 nM final siRNA concentration in each well.

The screening was performed in duplicate. The first replicate was analyzed in a luciferase assay while the latter was used for high-content imaging to enable visual inspection of the transfected cells. For luciferase measurements, plates were equilibrated for 10 min at room temperature and each well was aspirated to 10 µl volume. 12.5 µl of ONE-Glo reagent was added to each well and the plate was incubated at room temperature for 30 min. Luminescence was measured using the Safire2 (BioTek) microplate reader with 100 ms integration time. For the high-content imaging, the second replica plate was processed as in the first replicate and fixed using 35 µl/well 4% PFA for 30 min at room temperature. Cells were stained with 10 µM Hoechst for 1 h at room temperature. Plates were sealed and scanned using the ArrayScan™ XTI High Content Platform (Thermo Fisher).

### Drug Treatments

In the screening assays, cells were treated with the indicated agents for 4.5 h (25 µM Etoposide and 10 ng/ml TNFα). For the other experiments, unless otherwise stated, cells were treated with the following agents, at the indicated concentrations, and for the indicated amount of time: Etoposide (50 µM) for 90 min, TNFα (10 ng/ml) for 20 min, or olaparib (10 µM) for 12 h. Irradiation (20 Gy) was applied to cells by using a 137-Cs source (STS OB 29 device) 45 or 90 min, or at days indicated before experimental analysis.

### ONE-GLO^TM^+Tox Luciferase Reporter Assay

The luciferase assays were performed using a ONE-GLO^TM^+Tox kit according to the manufacturer’s protocol. In brief, 7000 cells (NF-κB/293/GFP-Luc^TM^) were seeded into 96 well plates with a black bottom and walls using a 100 µl growth medium RPMI supplemented with 5% penicillin/streptomycin (P/S) and 10% FBS, treated and transfected with indicated siRNAs if needed. The GF-FC dye was mixed in TOX substrate buffer in a 1:200 ratio and 20 µl of this solution was added to each well and incubated for 30 min at 37°C. The TOX cell viability signal was measured using a plate reader (Cytation 1) with 380 nm (Excitation) and 500 nm (Emission) filter sets. Subsequently, 100 µl of the luciferase substrate was added to each well and incubated for 3 min at RT. The luminescence signal was measured with the plate reader and normalized to the respective viability signal (TOX).

### siRNA and Plasmid Transfections

The transient knockdown of indicated genes was performed using Lipofectamine RNAiMAX according to the manufacturer’s protocols. In brief, the day after seeding cells into a 6 cm dish, at 2.5×10^5^ cells per dish quantity, 25 nM siRNA was mixed with 5 µl lipofectamine in 500 µl serum-free OptiMEM and incubated for 5 min at RT before adding to the cells in a drop-wise manner. The downstream experiments were performed 3 days after the transfection. A scrambled siRNA was used as a control for each experiment.

Ectopic expression of indicated recombinant DNA for the live-cell imaging experiments was performed using Lipofectamine 2000 according to the manufacturer’s protocols. In brief, the day after seeding cells into µ-Slide 8 well chamber (Ibidi), at 2.5×10^4^ cells per dish quantity, 1.5 µg recombinant DNA was mixed with 3 µl of lipofectamine in 50 µl serum-free OptiMEM and incubated for 5 min at RT before adding to the cells in a dropwise manner. For immunoprecipitation experiments, the indicated recombinant DNAs were transfected to the cells using polyethyleneimine (PEI). Cells were grown in 15 cm dishes at roughly 70% confluency, 15 µg of recombinant DNA was mixed with 90 µl of PEI in 2,85 ml serum and antibiotic-free DMEM and incubated for 30 min at RT before adding the cells. The downstream experiments were performed 2 days after the transfection.

### Immunoprecipitation

Immunoprecipitation of proteins was performed as previously described (Mikuda et al., 2018). In brief, cells were harvested in ice-cold CHAPS buffer (Tris-HCl pH 7.4, 110 mM NaCl, and 50 mM EDTA, supplemented with 1% (v/v) CHAPS, complete protease and phosphatase inhibitors cocktails), incubated with primary antibodies for overnight at 4°C, with G Fast Flow Sepharose Beads for 1 h at 4°C. Samples were boiled and subjected to standard SDS-PAGE.

Immunoprecipitation of GFP-tagged proteins was performed using GFP-trap (Chromotek) according to the manufacturer ‘s instructions. Briefly, cells were harvested in ice-cold RIPA buffer. Volume-matched 1 µg lysates were incubated with the bead slurry by end-over-end rotation for 1 hour at 4 °C. The beads were then sedimented and washed 3 times with RIPA buffer. After the last washing step, samples were mixed with 2X SDS-sample buffer and boiled. Beads were then sedimented again and the supernatants were subjected to standard SDS-PAGE.

### Protein Purification

293T cells were transfected with GFP-TSG101 plasmid using PEI. Cells were harvested in ice-cold RIPA buffer 48 h after transfection. GFP-TSG101 was immunoprecipitated using the GFP-trap method. To dissociate TSG101 from its binding partners, beads were additionally washed with high-salt (2M NaCl) and high detergent (1% Triton^TM^ X-100) containing wash buffers separately. Each washing step was repeated 3 times. Proteins were eluted from the beads using acidic elution buffer at room temperature. Solution was immediately neutralized with a neutralization buffer prior to storage.

### *In vitro* ADP Ribosylation Assay

*In vitro* ADP ribosylation was performed as previously described (Di Giammartino et al., 2013). The indicated amounts of TSG101 and PARP1 were incubated in buffer D (dialysis buffer) with 1mM NAD^+^, 400 ng sss (sheared salmon sperm) DNA, and 10 mM MgCl_2_. The reaction was carried out at 37 C° for 10 minutes and stopped by adding 2X SDS-loading buffer. After boiling, samples were subjected to standard SDS-PAGE.

### Total NAD and NADH Measurements

Total NAD and NADH measurements were performed using a colorimetric NAD/NADH assay kit according to manufacturer’s instructions. Briefly, 2.5×10^5^ cells were used for each condition. Cells were washed and scraped using ice-cold PBS. Cell pellets were mixed with NADH extraction buffer and the lysates were obtained by repeated freezing and thawing cycles. To remove any NADH consuming enzymes, lysates were filtered. For the detection of NADH, lysates were incubated at 60°C for 30 minutes. Under these conditions, all NAD^+^ will be decomposed while the NADH will still be intact. Samples were then cooled on ice. The NAD decomposed samples were converted to NADH using NAD Cycling Enzyme Mix. NADH developer was then added to the reaction and OD 450 nm reads were obtained using a plate reader.

Total NAD was quantified as in NADH measurement and NAD^+^ was calculated by subtracting NADH from the total NAD value.

### Cleaved Caspase-3 Measurement (Colorimetric)

Enzymatic activity of the cleaved caspase-3 was measured using a Caspase-3 Assay Kit (Colorimetric) (Abcam, ab39401), according to manufacturer’s instructions. Briefly, cells were harvested in PBS and cytoplasmic extracts were obtained. Protein concentrations were determined by a Bradford Assay and equal quantities of protein (100 µg for each condition) were tested in a colorimetric assay. Cleaved caspase-3 activity was measured using a plate-reader. Values were subtracted from an only buffer containing reaction and normalized to the level of non-targeting CRISPR transduced cells.

### CellTiter-Fluor^TM^ Cell Viability Assay

Cell viability was assayed using the CellTiter-Fluor^TM^ kit (Promega, G6081) according to manufacturer’s instructions. Briefly, cells were seeded onto 96-well plates with white walls and bottom at 10^5^ cells/well density. Indicated treatments and transfections were all performed in that plate. For cell viability measurements, GF-AFC substrate were added to each well. Cells were incubated at 37° for 30 min and cell viability was measured using a plate reader.

After subtracting the measurement from the reaction containing only medium, the values were normalized to the level of control siRNA transfected cells

### Western Blotting

For western blots, cell pellets were lysed in whole-cell extraction buffer (150 mM NaCl, 50 mM Tris pH 7.5, 1% NP-40 supplemented with proteinase and phosphatase inhibitors) for 20 min at 4°C while vigorously shaking, insoluble material was spun out, relative protein concentration was determined by BCA (BioRad). SDS sample buffer with 100 mM DTT was added to samples, which were loaded onto 8% Protein gels and separated by electrophoresis at 90-140 V for 2-3 h. Proteins were transferred and immobilized onto a nitrocellulose membrane (GE Healthcare) by electrophoresis for 2 h at RT in a standard transfer buffer containing 20% methanol. Membranes were blocked in 5% non-fat dry milk in TBS-T. All rabbit primary antibodies were probed with Peroxidase AffiniPure Donkey anti-rabbit IgG (Jackson ImmunoResearch) and all mouse primary antibodies were probed with Peroxidase AffiniPure Donkey anti-mouse IgG (Jackson ImmunoResearch). Chemiluminescent signals were visualized with Fusion Solo Imager.

### Proximity Ligation Assay (PLA)

The DuoLink (Merck) PLA assay was performed according to the manufacturer’s protocols. Cells were seeded in 12 mm glass coverslips 48 h before the experiment and treated if needed, fixed with 4% PFA at RT for 10 min, and permeabilized for 10 min at RT in PBS supplemented with 0.1% Triton X-100 (Sigma Aldrich). For blocking, cells were incubated for 1 h at 37°C in the provided blocking solution. Incubation with primary antibodies (PARP1, Thermo Fisher, Cat#MA 3-950, TSG101, Thermo Fisher, Cat#MA 1-23296, both diluted 1:200) was done overnight at 4°C.

### Laser Microirradiation and Live-cell Imaging

Cells were grown in µ-Slide 8 well chamber (Ibidi) and transiently transfected with GFP-tagged PARP1 and imaged at 24 h post-transfection. Throughout the experiment, cells were maintained in DMEM Dulbecco’s modified Eagle’s medium (DMEM) supplemented with and 10% FBS, at 37°C with 5% CO_2_. For sensitization, cells were incubated with a DNA damage sensitizer DAPI (10 µg/ml) for 10 min before the microirradiation. Cells with low to moderate GFP intensities were selected for the experiments. Targeted laser microirradition was induced by a 405 nm laser (100mW) using FRAPPA of the confocal spinning disk microscope (Andor CSU-W1 with Borealis on a Nikon Ti Eclipse microscope equipped with an iXON DU888 EMCCD camera). The laser power output was frequently measured to ensure constant DNA damage effect. A preselected area in the nucleus (a round-shaped with approximately 130 µm diameter) was microirraidated using a 100X CFI PLAN-Apo NA 1.4 oil-objective. The stimulation was performed using 25% laser power for 500 µs dwell time. In preliminary experiments the dosage 405 nm laser light was adjusted to the lowest possible amount of laser needed to induced DNA damage without compromising cell viability. Subsequent image acquisition was performed using a 488 nm laser line the same objective and an BP525/45 emission filter. Z-stacks of 20 slices were selected towards their top and bottom ends. The recruitment and dissociation of PARP1 to and from the DNA damage sites were followed with the time series up to 15 min with the first time point immediately after the micro-irradiation event.

For endogenous protein recruitment assays, cells were grown in µ-Slide 8 well-gridded chamber (Ibidi). The sensitization and transfection of the cells were done as in the live-cell imaging. To facilitate easy recognition of laser microirradiated cells during the imaging of endogenous TSG101, the DNA damage sites were created on a straight line in the nucleus of the selected cells but the parameters (405 nm laser wavelength with 25% laser power and 500 µs Dwell time) were kept consistent. Recruitment of PARP1 to the DNA damage site was recorded for either 1 or 5 min and cells were fixed with PFA and stained for TSG101 with standard indirect immunofluorescence. The microirradiated cells were refound and imaged with the confocal microscope (CSU Yokogawa Spinning Disk Field Scanning Confocal System, Nikon).

### Indirect Immunofluorescence

For all IF experiments (except the endogenous recruitment assay), cells were grown on 12 mm glass coverslips, treated if required, fixed in 4% PFA for 10 min at RT, and permeabilized for 10 min at RT in PBS supplemented with 0.1% Triton X-100 (Sigma Aldrich). Cells were blocked in filtered PBS supplemented with 0.1% Triton X-100 and 5% BSA for 1 h. The primary antibodies were diluted in the blocking buffer and cells were incubated with the antibody solution overnight at 4°C. All secondary antibodies were diluted in the blocking buffer and incubated for 1 h at RT. After each antibody incubation cells were washed 3 times with PBS. Following the last washing, coverslips were mounted with ProLong^TM^ Gold Antifade Mountant with DAPI. The primary antibodies used for IF at the indicated concentrations were against cleaved caspase 3 (Cell Signaling, 1:400), p65 (Cell Signaling, 1:500), PARP1 (Cell Signaling, 1:1000), and TSG101 (Thermo Fisher, 1:500). See Key Resources Table for the secondary antibodies and the fluorophores. Confocal immunofluorescence microscopy was done as previously described (Kolesnichenko et al., 2021; Mikuda et al., 2018).

### QRT-PCR

qRT-PCR was performed as previously described (Mikuda et al., 2018). In brief, primers were designed according to MIQE guidelines (Minimum Information for Publication of qRT-PCR Experiments) using NCBI primer blast and 62°C melting temperature was selected. To determine the fold induction of target genes, two to three reference genes (*ACTA1*, *RPL13A*, and *TBP2*) with an m value lower than 0.5 were used. The indicated primers were obtained from previous studies (Kolesnichenko et al., 2021; Mikuda et al., 2020; von Hoff et al., 2019). See Key Resources Table for qRT-PCR primers for details.

### Nuclear and Cytoplasmic Fractionation

The subcellular fractionation was performed as previously described (Mikuda et al., 2018). In brief, cells were washed with ice-cold PBS and harvested by scraping in 500 µl buffer A (10 mM Tris-HCl (pH 7.9, 1.5 mM MgCl_2_, 10 mM KCl, supplemented with complete protease and phosphatase inhibitors). After incubation on ice, NP-40 was added to a final concentration of 0.15% (v/v). Samples were washed, centrifuged and supernatants containing the cytoplasmic fraction were collected. The remaining pellet was further washed with buffer A containing NP-40 and resuspended two volumes of buffer C (20 mM Tris-HCl pH 7.9, 25% glycerol, 0.42 M NaCl, 1.5 mM MgCl_2_, 0.5% NP-40, supplemented with complete protease and phosphatase inhibitors). The insoluble nuclear debris was pelleted by centrifugation and nuclear lysates were subsequently collected.

### Electrophoretic Mobility Shift Assay (EMSA)

Nuclear extracts were used for the EMSA experiments. The assay was performed as previously described (Hinz et al., 2010). In brief, 5 µg of nuclear extract was mixed with radioactively labeled (P^32^) H2K probes supplemented with shift buffer (40 mM HEPES pH 7.9, 120 mM KCl, and 8% (v/v) Ficoll), 2 µg/ml poly dI/dC, 100 mM DTT and 10 µg/ml BSA and incubated for 30 min at RT. Samples were separated by electrophoresis and transferred to Whatman filter paper by vacuum dryer. The dried gel was incubated with Hyperfilm^TM^ MP at −80°C overnight and developed with standard procedures.

### CRISPR Cas9 knock-out Bulk Cell Generation

CRISPR Cas9-mediated knockout experiments were performed as previously described (Ran et al., 2013). In brief, the indicated guide RNA sequences were annealed and subcloned into lentiCRISPRv2 (Addgene, Cat#52961). The cloning was verified by Sanger sequencing. For lentiviral production, 293T cells were seeded into 15 cm plates at 90% confluency 1 day before transfection. The lentiCRISPRv2 vectors (30 µg) containing respective guide RNAs were transfected to the cells together with 20 µg of psPAX2 (Addgene, Cat#8454) and 10 µg of pCMV-VSV-G (Addgene, Cat#12260) using PEI. Lentiviruses were harvested and concentrated with PEG Virus Precipitation Kit (Abcam, Cat#ab102538) according to the manufacturer’s protocol. The target cells were seeded into 24-well plates at 20-40 % confluency and lentivirally transduced with the concentrated viruses at a 1:25 ratio in antibiotic-free DMEM supplemented with 10 µg/ml Polybrene (Sigma Aldrich). For the elimination of untransduced cells, puromycin (2 µg/ml) was added to the growth medium 3 days after the transduction. Knockout efficiencies were determined with western blotting. Due to the conclusive roles of TSG101 in cell division and development, a double knockout cell line was not established after single-cell expansion experiments.

### Crystal Violet Staining

Cells were seeded into 6-well plates, transfected, and treated if needed. The growth medium was gently aspirated and cells were fixed in ice-cold methanol for 10 min at 4°C. The plate was equilibrated to RT and incubated with crystal violet staining solution for 10 min at RT. The staining solution was removed and cells were washed with water. Crystal violet intensities were measured at 570 nm with a plate reader (Cytation 1, Biotek).

### Analysis of siRNA screening data

The statistical effect size (Z-factor) was calculated for each plate (189 in total) using the mock-transfected vehicle (DMSO) or etoposide (Sigma) treated cells negative and positive control respectively. The Z-factor cut-off was 0.1, 107 plates have a Z-factor below 0.5, 82 plates have a Z-factor equal or larger 0.5, the mean Z-Prime is 0.45. Several plates were repeated to improve quality. See Supplementary Table 2 for detailed Z-factor determination.

To determine assay reproducibility of the screening, 12 plates from the Druggable Genome Library were randomly selected for a replicate experiment. Results obtained from both replicates were compared and their correlation was determined by Spearman’s rank correlation coefficiency test (Supplementary Table 4).

For plate-wise normalization, the data obtained in the genome-wide screen was processed on the KNIME Analytics Platform (KNIME). A Z-score was calculated from luminescence values as follows: *x.zscore = (x – median(x[subset]))/mad(x[subset])*, where x is the luminescence value and the subset are all sample wells in the plate.

For the gene set enrichment analysis of the genome-wide screening hits, the REACTOME database (Jassal et al., 2020) was used.

### Analysis of quantitative live-cell imaging

Imaris software (Oxford Instruments) was used to quantify recruitment and dissociation kinetics of PARP1 to and from the DNA lesions. Images of pre and post-microirradiated cells were combined and the laser microirradiated area was segmented as a region of interest (ROI) in Imaris. The mean fluorescence intensity of the ROI was acquired for each time point. The fluorescence intensity of the whole nucleus was also acquired as a reference ROI and fluorescence intensity at the DNA damage site was normalized to the respective reference ROI.

### Statistical Tests

For statistical analysis, the normalized mean fluorescence intensities from 9 independent experiments were compared using an ordinary one-way ANOVA (ns, p > 0.05; *p < 0.05; **p < 0.01, ***p < 0.001; ****p < 0.0001).

## LEGENDS TO EXPANDED VIEW TABLES

**Expanded View Table 1: Z-score ranking of hits in the genome-wide siRNA screen**

Hits from the genome-wide siRNA screen were ranked according to their Z-scores (column D) indicating how far a given data point (relative luciferase units of the respective gene product) deviates from the average value of the complete dataset. Column A lists the plate IDs. DRUG indicates the part of the genome-wide siRNA library targeting druggable gene products.

**Expanded View Table 2: Z-Prime (Z-factor) determination of each plate used in the screening**

Z-Prime, which indicates separation between the distributions of the positive and negative controls and indicates the assay quality, was determined for each plate used in the screening. Plates are ranked according to their Z-Prime values. The Z-Prime cut-off is 0.1. The siRNA library consists of 189 plates. Several plates were repeated due to quality reasons and were accepted when the minimum Z-Prime criteria were met. 107 plates had a Z-Prime below 0.5, 82 plates had a Z-Prime equal or larger than 0.5, the mean Z-Prime was 0.45.

**Expanded View Table 3: Spearman correlation between randomly selected screening plates and their respective replicates**

For reproducibility assessment, 12 randomly selected plates were screened in duplicate. Respective replicates were compared using Spearman’s rank correlation coefficient test.

**Expanded View Table 4: Selected 320 genes from the genome-wide siRNA screen that were used for the TNFα counter screen**

320 candidates hits which led to decreased DNA damage-induced NF-κB activity in the genome-wide siRNA screen were selected for a counter screen with TNFα. Column A shows the gene names of the selected hits in alphabetical order.

**Expanded View Table 5: Luminescence values obtained for the 320 gene products and the respective controls in the TNFα counter screen.**

Column H shows NF-κB-driven luciferase activity of the respective hits. The first tab shows the etoposide-induced luminescence while the second tab shows the TNFα-induced luminescence as an NF-κB pathway readout. These results are visualized in Figure 1D.

**Expanded View Table 6: Hits from the “Metabolism of Lipids” category identified in the genome-wide siRNA screen.**

1000 candidate hits, which showed significantly decreased DNA damage-induced NF-κB activity in the genome-wide siRNA screen (putative positive regulators as indicated in Figure 1B) were analyzed in the REACTOME database (Jassal et al., 2020). Hits that are in the “Metabolism of Lipids” category are listed.

**Expanded View Table 7: Functional clustering of hits that positively regulate the NF-κB pathway. The hit selection section shows a comparison of the screening hits with other large-scale studies.**

Enrichment analysis of the putative positive regulatory hits. For each protein-coding gene, the keywords from the category “biological process” were derived from the UniProt knowledgebase and the abundance of these among the 1000 candidate hits and a comparison with the entire set of genes screened indicated the enrichment. The probability values (P-values) are listed in column H. The screening hits were compared with indicated large-scale studies and common hits are listed (Jungmichel et al., 2013; Mikuda et al., 2018; Paulsen et al., 2009).

**Expanded View Table 8: Materials used in this study**

List of used materials, including antibodies, chemicals, peptides, recombinant proteins, commercial assays, recombinant DNA and plasmids, oligonucleotides and software.

## LEGENDS TO EXPANDED VIEW FIGURES

**Expanded View Figure 1.**

A: HEK-Luc(tGFP) NF-κB reporter cells were treated with the indicated conditions (50 µM etoposide, 20 Gy irradiation, or 10 ng/ml TNFα). As the NF-κB-driven expression and translation of turbo GFP requires more time than the initial activation of the NF-κB pathway, the treatment conditions were alternatively prolonged to 4.5 hours. Whole-cell extracts were obtained and immunoblotted with the indicated antibodies.

B: Detailed layout of the genome-wide siRNA screening, carried out using a one-plate-to-one target approach. The indicated controls were added to the siRNA library. As another control, cells were treated with the vehicle DMSO (Column 24). The heatmap displays the Z-scores of tested candidates and controls of a representative plate taken from the siRNA library screen.

C: Spearman’s rank correlation coefficient test of representative screening plate replicates demonstrated high assay reproducibility. Same screening plates and layouts were tested and results compared with each other (R1 vs R2).

D: TSG101 cluster was visualized using STRING database (Szklarczyk et al., 2019). Arrows or lines between candidate hits represent protein-protein interactions observed in previous publications.

E: Representative high-confidence DNA damage-selective hits are shown for etoposide (left panel) and TNFα (right panel) screens. The conditions were compared with an ordinary one-way ANOVA (ns, p > 0.05; *p < 0.05; **p < 0.01, ***p < 0.001; ****p < 0.0001).

**Expanded View Figure 2.**

A: HEK-Luc(tGFP) NF-κB reporter cells were transfected with the same transfection mixtures as used in Fig 2A. Whole-cell extracts were immunoblotted with antibodies against the indicated proteins. Actin was used as a loading control.

B: U2-OS cells were transfected with the indicated siRNAs. The NF-ĸB pathway was activated by etoposide treatment (50 µM, 90 minutes before analysis). Expression of indicated genes was analyzed using qRT-PCR. The mRNA expression of these genes was normalized to the expression of three housekeeping genes, *ACTA1*, *RPL13A*, and *TBP2*. The gene expression for the indicated conditions are relative to the non-targeting siRNA-transfected vehicle (DMSO)-treated cells. The result is representative of three biologically independent experiments. The conditions were compared with an ordinary one-way ANOVA (ns, p > 0.05; *p < 0.05; **p < 0.01, ***p < 0.001; ****p < 0.0001).

**Expanded View Figure 3.**

A: U2-OS cells were transfected with non-targeting or *TSG101*-targeting siRNAs. NADH measurements were performed 72 hours after the siRNA transfection. Conditions were compared with Student’s t-test (Welch Correction).

B: U2-OS cells were transfected with non-targeting (Mock) or *TSG101*-directed siRNAs, with mCherry-*TSG101,* or treated with olaparib (10 µM, 16 h before irradiation), as indicated. Cells were irradiated (20 Gy) 10 min. before analysis and NAD+/NADH levels were determined. The result is representative of four independent experiments. The conditions were compared with an ordinary one-way ANOVA (ns, p > 0.05; *p < 0.05; **p < 0.01, ***p < 0.001; ****p < 0.0001).

C: U2-OS cells were transfected with non-targeting (-) or *ACADVL*-targeting (+) siRNAs. Cells were treated with etoposide (50 µM, 10 minutes before analysis), as indicated. Whole-cell extracts were immunoblotted with 10H PAR antibodies. Actin was used as a loading control. The result is representative of three biologically independent experiments.

D: U2-OS cells were transfected with non-targeting (-) or *NAA15*-targeting (+) siRNAs. Cells were treated with etoposide (50 µM, 10 minutes before analysis). Whole-cell extracts were immunoblotted with antibodies against the indicated proteins or posttranslational modifications. Vinculin was used as a loading control. The result is representative of three biologically independent experiments.

E and F: U2-OS cells were transfected with non-targeting (-) or two different each *ACADVL-* or *NAA15*-targeting [+, (1) or (2)] siRNAs. Cells were treated with etoposide (+) (50 µM, 90 minutes before analysis), or not (-). Whole-cell extracts were immunoblotted with antibodies against the indicated proteins or posttranslational modifications. Actin was used as a loading control. The results are representative of three biologically independent experiments.

G: U2-OS cells were transfected with non-targeting (-) or NAA15-targeting (+) siRNAs. Cells were irradiated (IR) (+, 20 Gy, 30 minutes before analysis) or not (-). Whole-cell extracts were immunoblotted with antibodies against the indicated proteins. Actin was used as a loading control. The results are representative of three biologically independent experiments.

**Expanded View Figure 4.**

A: Final protein-protein interaction probability scores for the indicated interactions were obtained from the PrePPI database (Zhang et al., 2013).

B: U2-OS cells were transfected with the full-length mCherry-tagged TSG101 plasmid and DNA damage was induced by irradiation (20 Gy, 45 min before analysis), as indicated. Immunoprecipitation of mCherry was performed using whole-cell extracts. The result is representative of three biologically independent experiments.

**Expanded View Figure 5.**

A: U2-OS cells were transfected with non-targeting (Mock) or *TSG101*-targeting siRNAs. Indirect immunofluorescence visualizes TSG101 (red) and PARP1 (green). DAPI staining shows the nuclei in blue. Scale bar is 10 µm. The image is representative of three biologically independent experiments.

B: Venn diagram showing the intersection of the positive regulators from the genome-wide screen and the ESCRT family proteins.

C: U2-OS cells were transfected with non-targeting or *UBAP1*-targeting siRNAs. Whole-cell extracts were immunoblotted with antibodies against the indicated proteins. Vinculin was used as a loading control.

D: Total RNA was extracted from cells analyzed in B and converted to cDNA. The relative mRNA expression of *UBAP1* was normalized to two housekeeping genes, *RPL13A*, and *TBP2.* Data are from six biologically independent experiments. The conditions were compared with an unpaired t test with Welch’s correction (ns, p > 0.05; *p < 0.05; **p < 0.01, ***p < 0.001; ****p < 0.0001).

**Expanded View Figure 6.**

A: U2-OS cells were lentivirally transduced with two independent guide RNAs targeting the *TSG101* locus. Due to the lethal side effects of TSG101 deletion on long-term proliferation, bulk cells were used instead of the clonally expanded cells. DNA damage was induced by irradiation (20 Gy, 3 or 9 hours before analysis) in wild type or TSG101 CRISPR knockout bulk cells. Total mRNA was extracted and expression of *NUAK*, *PTX3*, and *PUMA* was analyzed with RT-qPCR. The mRNA expression was normalized to the housekeeping genes *ACTA1*, *RPL13A*, and *TBP2*. Data are from three biologically independent experiments. The conditions were compared with an ordinary one-way ANOVA (ns, p > 0.05; *p < 0.05; **p < 0.01, ***p < 0.001; ****p < 0.0001).

B: The efficiency of *TSG101* CRISPR in the cells used in A is shown with immunoblotting. Whole-cell extracts were immunoblotted with antibodies against TSG101. Vinculin was used as a loading control.

C: Irradiation-induced cleaved caspase-3 activation in non-targeting gRNA or *Tsg101-*targeting gRNA transduced MEF cells was measured using a colorimetric assay. For knock-out efficiencies see Fig EV6E. Bulk cells from *Tsg101* guide-1 were used in this assay.

D: Percentage of γH2AX positive staining from Fig 6D is shown.

E: Wild-type MEF cells from Fig 6J (control or *Tsg101* guide RNA transduced cells) were analyzed for *Tsg101* CRSIPR/Cas9 knock-out efficiency. Whole-cell extracts were obtained from these cells and immunoblotted with the indicated antibodies. Actin was used as a loading control.

F: U2-OS cells were transfected with non-targeting or *TSG101*-targeting siRNAs. The 7 days post-irradiation time point was selected to analyze senescent cells. Total mRNA was extracted and expression of *CXCL8*, *IL-6*, and *TSG101* was analyzed by RT-qPCR. The mRNA expression of these genes was normalized to the three housekeeping genes *ACTA1*, *RPL13A* and *TBP2*. Data are from three biologically independent experiments. The conditions were compared with an ordinary one-way ANOVA (ns, p > 0.05; *p < 0.05; **p < 0.01, ***p < 0.001; ****p < 0.0001).

G: Representative brightfield images of *TSG101*-targeting or non-targeting siRNA transfected *BRCA1* wild type and mutant MDA-MB 231 and 436 breast cancer cells are shown.

## REFERENCES

Ahel, D., Horejsí, Z., Wiechens, N., Polo, S.E., Garcia-Wilson, E., Ahel, I., Flynn, H., Skehel, M., West, S.C., Jackson, S.P., Owen-Hughes, T., Boulton, S.J., 2009. Poly(ADP-ribose)-dependent regulation of DNA repair by the chromatin remodeling enzyme ALC1. Science 325, 1240– 1243. https://doi.org/10.1126/science.1177321

Arnesen, T., Anderson, D., Baldersheim, C., Lanotte, M., Varhaug, J.E., Lillehaug, J.R., 2005. Identification and characterization of the human ARD1-NATH protein acetyltransferase complex. Biochem J 386, 433–443. https://doi.org/10.1042/BJ20041071

Babst, M., Odorizzi, G., Estepa, E.J., Emr, S.D., 2000. Mammalian tumor susceptibility gene 101 (TSG101) and the yeast homologue, Vps23p, both function in late endosomal trafficking. Traffic 1, 248–258. https://doi.org/10.1034/j.1600-0854.2000.010307.x

Blessing, C., Mandemaker, I.K., Gonzalez-Leal, C., Preisser, J., Schomburg, A., Ladurner, A.G., 2020. The Oncogenic Helicase ALC1 Regulates PARP Inhibitor Potency by Trapping PARP2 at DNA Breaks. Mol Cell 80, 862–875.e6. https://doi.org/10.1016/j.molcel.2020.10.009

Bonner, W.M., Redon, C.E., Dickey, J.S., Nakamura, A.J., Sedelnikova, O.A., Solier, S., Pommier, Y., 2008. GammaH2AX and cancer. Nat Rev Cancer 8, 957–967. https://doi.org/10.1038/nrc2523

Callen, E., Di Virgilio, M., Kruhlak, M.J., Nieto-Soler, M., Wong, N., Chen, H.-T., Faryabi, R.B., Polato, F., Santos, M., Starnes, L.M., Wesemann, D.R., Lee, J.-E., Tubbs, A., Sleckman, B.P., Daniel, J.A., Ge, K., Alt, F.W., Fernandez-Capetillo, O., Nussenzweig, M.C., Nussenzweig, A., 2013. 53BP1 mediates productive and mutagenic DNA repair through distinct phosphoprotein interactions. Cell 153, 1266–1280. https://doi.org/10.1016/j.cell.2013.05.023

Colomer, C., Margalef, P., Villanueva, A., Vert, A., Pecharroman, I., Solé, L., González-Farré, M., Alonso, J., Montagut, C., Martinez-Iniesta, M., Bertran, J., Borràs, E., Iglesias, M., Sabidó, E., Bigas, A., Boulton, S.J., Espinosa, L., 2019. IKKα Kinase Regulates the DNA Damage Response and Drives Chemo-resistance in Cancer. Mol Cell 75, 669–682.e5. https://doi.org/10.1016/j.molcel.2019.05.036

Cortes, U., Tong, W.-M., Coyle, D.L., Meyer-Ficca, M.L., Meyer, R.G., Petrilli, V., Herceg, Z., Jacobson, E.L., Jacobson, M.K., Wang, Z.-Q., 2004. Depletion of the 110-kilodalton isoform of poly(ADP-ribose) glycohydrolase increases sensitivity to genotoxic and endotoxic stress in mice. Mol Cell Biol 24, 7163–7178. https://doi.org/10.1128/MCB.24.16.7163-7178.2004

D’Amours, D., Desnoyers, S., D’Silva, I., Poirier, G.G., 1999. Poly(ADP-ribosyl)ation reactions in the regulation of nuclear functions. Biochem J 342 (Pt 2), 249–268.

Di Giammartino, D.C., Shi, Y., Manley, J.L., 2013. PARP1 represses PAP and inhibits polyadenylation during heat shock. Mol Cell 49, 7–17. https://doi.org/10.1016/j.molcel.2012.11.005

Doyotte, A., Russell, M.R.G., Hopkins, C.R., Woodman, P.G., 2005. Depletion of TSG101 forms a mammalian “Class E” compartment: a multicisternal early endosome with multiple sorting defects. J Cell Sci 118, 3003–3017. https://doi.org/10.1242/jcs.02421

Dunphy, G., Flannery, S.M., Almine, J.F., Connolly, D.J., Paulus, C., Jønsson, K.L., Jakobsen, M.R., Nevels, M.M., Bowie, A.G., Unterholzner, L., 2018. Non-canonical Activation of the DNA Sensing Adaptor STING by ATM and IFI16 Mediates NF-κB Signaling after Nuclear DNA Damage. Mol Cell 71, 745–760.e5. https://doi.org/10.1016/j.molcel.2018.07.034

Ferraiuolo, R.-M., Manthey, K.C., Stanton, M.J., Triplett, A.A., Wagner, K.-U., 2020. The Multifaceted Roles of the Tumor Susceptibility Gene 101 (TSG101) in Normal Development and Disease. Cancers (Basel) 12. https://doi.org/10.3390/cancers12020450

Fong, P.C., Boss, D.S., Yap, T.A., Tutt, A., Wu, P., Mergui-Roelvink, M., Mortimer, P., Swaisland, H., Lau, A., O’Connor, M.J., Ashworth, A., Carmichael, J., Kaye, S.B., Schellens, J.H.M., de Bono, J.S., 2009. Inhibition of poly(ADP-ribose) polymerase in tumors from BRCA mutation carriers. N Engl J Med 361, 123–134. https://doi.org/10.1056/NEJMoa0900212

Fu, J., Huang, D., Yuan, F., Xie, N., Li, Q., Sun, X., Zhou, X., Li, G., Tong, T., Zhang, Y., 2018. TRAF-interacting protein with forkhead-associated domain (TIFA) transduces DNA damage–induced activation of NF-κB. J Biol Chem 293, 7268–7280. https://doi.org/10.1074/jbc.RA117.001684

Genois, M.-M., Gagné, J.-P., Yasuhara, T., Jackson, J., Saxena, S., Langelier, M.-F., Ahel, I., Bedford, M.T., Pascal, J.M., Vindigni, A., Poirier, G.G., Zou, L., 2021. CARM1 regulates replication fork speed and stress response by stimulating PARP1. Mol Cell 81, 784–800.e8. https://doi.org/10.1016/j.molcel.2020.12.010

Gibbs-Seymour, I., Fontana, P., Rack, J.G.M., Ahel, I., 2016. HPF1/C4orf27 Is a PARP-1-Interacting Protein that Regulates PARP-1 ADP-Ribosylation Activity. Mol Cell 62, 432–442. https://doi.org/10.1016/j.molcel.2016.03.008

Gibson, B.A., Kraus, W.L., 2012. New insights into the molecular and cellular functions of poly(ADP-ribose) and PARPs. Nature Reviews Molecular Cell Biology 13, 411–424. https://doi.org/10.1038/nrm3376

Goldstein, M., Kastan, M.B., 2015. The DNA damage response: implications for tumor responses to radiation and chemotherapy. Annu Rev Med 66, 129–143. https://doi.org/10.1146/annurev-med-081313-121208

Haskill, S., Beg, A.A., Tompkins, S.M., Morris, J.S., Yurochko, A.D., Sampson-Johannes, A., Mondal, K., Ralph, P., Baldwin, A.S., 1991. Characterization of an immediate-early gene induced in adherent monocytes that encodes I kappa B-like activity. Cell 65, 1281–1289. https://doi.org/10.1016/0092-8674(91)90022-q

Helleday, T., 2011. The underlying mechanism for the PARP and BRCA synthetic lethality: Clearing up the misunderstandings. Molecular Oncology, Genetic Instability and Cancer 5, 387–393. https://doi.org/10.1016/j.molonc.2011.07.001

Hinz, M., Stilmann, M., Arslan, S.Ç., Khanna, K.K., Dittmar, G., Scheidereit, C., 2010. A cytoplasmic ATM-TRAF6-cIAP1 module links nuclear DNA damage signaling to ubiquitin-mediated NF-κB activation. Mol Cell 40, 63–74. https://doi.org/10.1016/j.molcel.2010.09.008

Hopp, A.-K., Teloni, F., Bisceglie, L., Gondrand, C., Raith, F., Nowak, K., Muskalla, L., Howald, A., Pedrioli, P.G.A., Johnsson, K., Altmeyer, M., Pedrioli, D.M.L., Hottiger, M.O., 2021. Mitochondrial NAD+ Controls Nuclear ARTD1-Induced ADP-Ribosylation. Mol Cell 81, 340–354.e5. https://doi.org/10.1016/j.molcel.2020.12.034

Huang, T.T., Wuerzberger-Davis, S.M., Wu, Z.-H., Miyamoto, S., 2003. Sequential modification of NEMO/IKKgamma by SUMO-1 and ubiquitin mediates NF-kappaB activation by genotoxic stress. Cell 115, 565–576. https://doi.org/10.1016/s0092-8674(03)00895-x

Jackson, S.P., Bartek, J., 2009. The DNA-damage response in human biology and disease. Nature 461, 1071–1078. https://doi.org/10.1038/nature08467

Jassal, B., Matthews, L., Viteri, G., Gong, C., Lorente, P., Fabregat, A., Sidiropoulos, K., Cook, J., Gillespie, M., Haw, R., Loney, F., May, B., Milacic, M., Rothfels, K., Sevilla, C., Shamovsky, V., Shorser, S., Varusai, T., Weiser, J., Wu, G., Stein, L., Hermjakob, H., D’Eustachio, P., 2020. The reactome pathway knowledgebase. Nucleic Acids Res 48, D498–D503. https://doi.org/10.1093/nar/gkz1031

Jungmichel, S., Rosenthal, F., Altmeyer, M., Lukas, J., Hottiger, M.O., Nielsen, M.L., 2013. Proteome-wide identification of poly(ADP-Ribosyl)ation targets in different genotoxic stress responses. Mol Cell 52, 272–285. https://doi.org/10.1016/j.molcel.2013.08.026

Kim, M.Y., Zhang, T., Kraus, W.L., 2005. Poly(ADP-ribosyl)ation by PARP-1: “PAR-laying” NAD+ into a nuclear signal. Genes Dev 19, 1951–1967. https://doi.org/10.1101/gad.1331805

Kolesnichenko, M., Mikuda, N., Höpken, U.E., Kärgel, E., Uyar, B., Tufan, A.B., Milanovic, M., Sun, W., Krahn, I., Schleich, K., von Hoff, L., Hinz, M., Willenbrock, M., Jungmann, S., Akalin, A., Lee, S., Schmidt-Ullrich, R., Schmitt, C.A., Scheidereit, C., 2021. Transcriptional repression of NFKBIA triggers constitutive IKK-and proteasome-independent p65/RelA activation in senescence. The EMBO Journal 40, e104296. https://doi.org/10.15252/embj.2019104296

Krishnakumar, R., Kraus, W.L., 2010. The PARP side of the nucleus: molecular actions, physiological outcomes, and clinical targets. Mol Cell 39, 8–24. https://doi.org/10.1016/j.molcel.2010.06.017

Kuleshov, M.V., Jones, M.R., Rouillard, A.D., Fernandez, N.F., Duan, Q., Wang, Z., Koplev, S., Jenkins, S.L., Jagodnik, K.M., Lachmann, A., McDermott, M.G., Monteiro, C.D., Gundersen, G.W., Ma’ayan, A., 2016. Enrichr: a comprehensive gene set enrichment analysis web server 2016 update. Nucleic Acids Res 44, W90–97. https://doi.org/10.1093/nar/gkw377

Lee, M.H., Mabb, A.M., Gill, G.B., Yeh, E.T.H., Miyamoto, S., 2011. NF-κB induction of the SUMO protease SENP2: A negative feedback loop to attenuate cell survival response to genotoxic stress. Mol Cell 43, 180–191. https://doi.org/10.1016/j.molcel.2011.06.017

Li, N., Banin, S., Ouyang, H., Li, G.C., Courtois, G., Shiloh, Y., Karin, M., Rotman, G., 2001. ATM is required for IkappaB kinase (IKKk) activation in response to DNA double strand breaks. J Biol Chem 276, 8898–8903. https://doi.org/10.1074/jbc.M009809200

Luo, X., Kraus, W.L., 2012. On PAR with PARP: cellular stress signaling through poly(ADP-ribose) and PARP-1. Genes Dev 26, 417–432. https://doi.org/10.1101/gad.183509.111

Mabb, A.M., Wuerzberger-Davis, S.M., Miyamoto, S., 2006. PIASy mediates NEMO sumoylation and NF-κB activation in response to genotoxic stress. Nature Cell Biology 8, 986–993. https://doi.org/10.1038/ncb1458

Mamińska, A., Bartosik, A., Banach-Orłowska, M., Pilecka, I., Jastrzębski, K., Zdżalik-Bielecka, D., Castanon, I., Poulain, M., Neyen, C., Wolińska-Nizioł, L., Toruń, A., Szymańska, E., Kowalczyk, A., Piwocka, K., Simonsen, A., Stenmark, H., Fürthauer, M., González-Gaitán, M., Miaczynska, M., 2016. ESCRT proteins restrict constitutive NF-κB signaling by trafficking cytokine receptors. Sci Signal 9, ra8. https://doi.org/10.1126/scisignal.aad0848

Maya-Mendoza, A., Moudry, P., Merchut-Maya, J.M., Lee, M., Strauss, R., Bartek, J., 2018. High speed of fork progression induces DNA replication stress and genomic instability. Nature 559, 279– 284. https://doi.org/10.1038/s41586-018-0261-5

McAndrew, R.P., Wang, Y., Mohsen, A.-W., He, M., Vockley, J., Kim, J.-J.P., 2008. Structural basis for substrate fatty acyl chain specificity: crystal structure of human very-long-chain acyl-CoA dehydrogenase. J Biol Chem 283, 9435–9443. https://doi.org/10.1074/jbc.M709135200

Mikuda, N., Kolesnichenko, M., Beaudette, P., Popp, O., Uyar, B., Sun, W., Tufan, A.B., Perder, B., Akalin, A., Chen, W., Mertins, P., Dittmar, G., Hinz, M., Scheidereit, C., 2018. The IκB kinase complex is a regulator of mRNA stability. The EMBO Journal 37, e98658. https://doi.org/10.15252/embj.201798658

Mikuda, N., Schmidt-Ullrich, R., Kärgel, E., Golusda, L., Wolf, J., Höpken, U.E., Scheidereit, C., Kühl, A.A., Kolesnichenko, M., 2020. Deficiency in IκBα in the intestinal epithelium leads to spontaneous inflammation and mediates apoptosis in the gut. J Pathol 251, 160–174. https://doi.org/10.1002/path.5437

Mortusewicz, O., Amé, J.-C., Schreiber, V., Leonhardt, H., 2007. Feedback-regulated poly(ADP-ribosyl)ation by PARP-1 is required for rapid response to DNA damage in living cells. Nucleic Acids Res 35, 7665–7675. https://doi.org/10.1093/nar/gkm933

Mueller, F., Friese, A., Pathe, C., da Silva, R.C., Rodriguez, K.B., Musacchio, A., Bange, T., 2021. Overlap of NatA and IAP substrates implicates N-terminal acetylation in protein stabilization. Sci Adv 7, eabc8590. https://doi.org/10.1126/sciadv.abc8590

Piret, B., Schoonbroodt, S., Piette, J., 1999. The ATM protein is required for sustained activation of NF-kappaB following DNA damage. Oncogene 18, 2261–2271. https://doi.org/10.1038/sj.onc.1202541

Polo, S.E., Jackson, S.P., 2011. Dynamics of DNA damage response proteins at DNA breaks: a focus on protein modifications. Genes Dev 25, 409–433. https://doi.org/10.1101/gad.2021311

Ran, F.A., Hsu, P.D., Wright, J., Agarwala, V., Scott, D.A., Zhang, F., 2013. Genome engineering using the CRISPR-Cas9 system. Nat Protoc 8, 2281–2308. https://doi.org/10.1038/nprot.2013.143

Ruland, J., Sirard, C., Elia, A., MacPherson, D., Wakeham, A., Li, L., Luis de la Pompa, J., Cohen, S.N., Mak, T.W., 2001. p53 Accumulation, defective cell proliferation, and early embryonic lethality in mice lacking tsg101. Proc Natl Acad Sci U S A 98, 1859–1864.

Shao, L., Zhou, H.J., Zhang, H., Qin, L., Hwa, J., Yun, Z., Ji, W., Min, W., 2015. SENP1-mediated NEMO deSUMOylation in adipocytes limits inflammatory responses and type-1 diabetes progression. Nat Commun 6, 8917. https://doi.org/10.1038/ncomms9917

Shi, H., Sun, L., Wang, Y., Liu, A., Zhan, X., Li, X., Tang, M., Anderton, P., Hildebrand, S., Quan, J., Ludwig, S., Moresco, E.M.Y., Beutler, B., 2021. N4BP1 negatively regulates NF-κB by binding and inhibiting NEMO oligomerization. Nat Commun 12, 1379. https://doi.org/10.1038/s41467-021-21711-5

Solier, S., Pommier, Y., 2009. The apoptotic ring: a novel entity with phosphorylated histones H2AX and H2B and activated DNA damage response kinases. Cell Cycle 8, 1853–1859. https://doi.org/10.4161/cc.8.12.8865

Stilmann, M., Hinz, M., Arslan, S.C., Zimmer, A., Schreiber, V., Scheidereit, C., 2009. A nuclear poly(ADP-ribose)-dependent signalosome confers DNA damage-induced IkappaB kinase activation. Mol Cell 36, 365–378. https://doi.org/10.1016/j.molcel.2009.09.032

Sun, S.-C., 2017. The non-canonical NF-κB pathway in immunity and inflammation. Nat Rev Immunol 17, 545–558. https://doi.org/10.1038/nri.2017.52

Szklarczyk, D., Gable, A.L., Lyon, D., Junge, A., Wyder, S., Huerta-Cepas, J., Simonovic, M., Doncheva, N.T., Morris, J.H., Bork, P., Jensen, L.J., Mering, C. von, 2019. STRING v11: protein-protein association networks with increased coverage, supporting functional discovery in genome-wide experimental datasets. Nucleic Acids Res 47, D607–D613. https://doi.org/10.1093/nar/gky1131

Thul, P.J., Lindskog, C., 2018. The human protein atlas: A spatial map of the human proteome. Protein Sci 27, 233–244. https://doi.org/10.1002/pro.3307

Trompouki, E., Hatzivassiliou, E., Tsichritzis, T., Farmer, H., Ashworth, A., Mosialos, G., 2003. CYLD is a deubiquitinating enzyme that negatively regulates NF-kappaB activation by TNFR family members. Nature 424, 793–796. https://doi.org/10.1038/nature01803

Tsherniak, A., Vazquez, F., Montgomery, P.G., Weir, B.A., Kryukov, G., Cowley, G.S., Gill, S., Harrington, W.F., Pantel, S., Krill-Burger, J.M., Meyers, R.M., Ali, L., Goodale, A., Lee, Y., Jiang, G., Hsiao, J., Gerath, W.F.J., Howell, S., Merkel, E., Ghandi, M., Garraway, L.A., Root, D.E., Golub, T.R., Boehm, J.S., Hahn, W.C., 2017. Defining a Cancer Dependency Map. Cell 170, 564–576.e16. https://doi.org/10.1016/j.cell.2017.06.010

Vallabhapurapu, S., Matsuzawa, A., Zhang, W., Tseng, P.-H., Keats, J.J., Wang, H., Vignali, D.A.A., Bergsagel, P.L., Karin, M., 2008. Nonredundant and complementary functions of TRAF2 and TRAF3 in a ubiquitination cascade that activates NIK-dependent alternative NF-kappaB signaling. Nat Immunol 9, 1364–1370. https://doi.org/10.1038/ni.1678

Vietri, M., Radulovic, M., Stenmark, H., 2020. The many functions of ESCRTs. Nat Rev Mol Cell Biol 21, 25–42. https://doi.org/10.1038/s41580-019-0177-4

von Hoff, L., Kärgel, E., Franke, V., McShane, E., Schulz-Beiss, K.W., Patone, G., Schleussner, N., Kolesnichenko, M., Hübner, N., Daumke, O., Selbach, M., Akalin, A., Mathas, S., Scheidereit, C., 2019. Autocrine LTA signaling drives NF-κB and JAK-STAT activity and myeloid gene expression in Hodgkin lymphoma. Blood 133, 1489–1494. https://doi.org/10.1182/blood-2018-08-871293

Wang, W., Huang, X., Xin, H.-B., Fu, M., Xue, A., Wu, Z.-H., 2015. TRAF Family Member-associated NF-κB Activator (TANK) Inhibits Genotoxic Nuclear Factor κB Activation by Facilitating Deubiquitinase USP10-dependent Deubiquitination of TRAF6 Ligase. J Biol Chem 290, 13372– 13385. https://doi.org/10.1074/jbc.M115.643767

Wang, W., Mani, A.M., Wu, Z.-H., 2017. DNA damage-induced nuclear factor-kappa B activation and its roles in cancer progression. J Cancer Metastasis Treat 3, 45–59. https://doi.org/10.20517/2394-4722.2017.03

White, J.T., Rives, J., Tharp, M.E., Wrabl, J.O., Thompson, E.B., Hilser, V.J., 2021. Tumor Susceptibility Gene 101 Regulates the Glucocorticoid Receptor through Disorder-Mediated Allostery. Biochemistry 60, 1647–1657. https://doi.org/10.1021/acs.biochem.1c00079

Wu, Z.-H., Shi, Y., Tibbetts, R.S., Miyamoto, S., 2006. Molecular linkage between the kinase ATM and NF-kappaB signaling in response to genotoxic stimuli. Science 311, 1141–1146. https://doi.org/10.1126/science.1121513

Zhang, Q.C., Petrey, D., Garzón, J.I., Deng, L., Honig, B., 2013. PrePPI: a structure-informed database of protein-protein interactions. Nucleic Acids Res 41, D828–833. https://doi.org/10.1093/nar/gks1231

